# Two roles for choice selective inhibition in decision-making circuits

**DOI:** 10.1101/2022.01.24.477635

**Authors:** James P. Roach, Anne K. Churchland, Tatiana A. Engel

## Abstract

During perceptual decision-making, the firing rates of cortical neurons reflect upcoming choices. Recent work showed that excitatory and inhibitory neurons are equally selective for choice. However, functional consequences of inhibitory choice selectivity in decision-making circuits are unknown. We developed a circuit model of decision-making which accounts for the specificity of inputs to and outputs from inhibitory neurons. We found that selective inhibition expands the space of circuits supporting decision-making, allowing for weaker or stronger recurrent excitation when connected in a competitive or feedback motif. The specificity of inhibitory outputs sets the trade-off between speed and accuracy of decisions by altering the attractor dynamics in the circuit. Recurrent neural networks trained to make decisions display the same dependence on inhibitory specificity and the strength of recurrent excitation. Our results reveal two concurrent roles for selective inhibition in decision-making circuits: stabilizing strongly connected excitatory populations and maximizing competition between oppositely selective populations.

---

Perceptual decision-making requires neural circuits to integrate evidence and classify a stimulus to trigger the correct behavioral response. Neurons in a range of cortical areas modulate their firing rate to signal animal’s choice^1^. The functional properties of decision-making neural circuits have been extensively studied and modeled^2–9^. Central to the function of these circuit models are attractors in the activity space which characterize the population’s encoding of a given choice. The attractor mechanism driving the decision-making activity in these models relies on highly structured recurrent connections between populations of excitatory neurons that are each selective for a different choice^8, 10, 11^. Inhibitory neurons, in this view, are merely supporting actors facilitating competition and providing balance to the excitatory neurons.

Since the canonical models of decision-making circuits were built, the diversity and complexity of inhibitory neurons within the cortex have been characterized in increasing detail^12^. In primary sensory areas, inhibitory neurons are generally more broadly tuned^13^ and more densely connected to neighboring excitatory neurons^14, 15^. These inhibitory neurons reliably modulate spike output to reflect stimulus features and have highly specific connectivity to surrounding excitatory neurons^16, 17^. The stimulus selectivity of inhibitory neurons is enhanced by learning and attention^18^ suggesting that task dependent modulation of inhibitory activity is necessary for cognition. Beyond the primary sensory cortex, stimulus information and animal choice can be decoded from the activity of inhibitory neurons in secondary sensory and association areas indicating a role for selective inhibition in higher cognitive functions, such as decision making^19–21^. While there is growing evidence that the activity and connectivity of inhibitory neurons is as complex as excitatory neurons, how the selectivity of inhibitory activity and the diversity of their connections affect the decision-making function of cortical circuits is still unknown.

To reveal the role of choice selective inhibitory neurons in decision-making computations we extended a well established mean-field model of decision-making circuits^4^ to account for the presence of inhibitory choice selectivity. Our model allows us to parametrically alter the specificity of connections between four choice selective populations: two excitatory and two inhibitory. Through analysis of this model, we identified two concurrent roles for inhibition in decision-making circuits: inhibition drives competition between choice-selective excitatory populations and at the same time stabilizes activity driven by recurrent excitation. These two roles are mediated by inhibitory connections to the excitatory populations and either role can be enhanced by structured inhibitory connectivity. We found that inhibitory selectivity expands the space of possible circuits which support decision-making by enhancing either a competitive or stabilizing role for inhibition. In addition, the connectivity motif between choice selective populations alters the underlying attractor dynamics and modulates the decision-making performance to prioritize speed or accuracy. We generalized these results by training recurrent neural networks (RNNs) to perform the same decision-making task. After training, RNNs had both excitatory and inhibitory units significantly selective for choice and displayed a similar dependence between the specificity of excitatory and inhibitory connections found in the mean-field model. Finally, we perturbed inhibitory neuron activity in these models to probe the dynamical regime in which the circuit operates. We found two regimes in which circuits respond differently to perturbations of inhibitory neurons: one in which the competitive role dominates and the other in which the stabilizing role dominates. Our work demonstrates that choice selective inhibition impacts decision-making behavior by enhancing either the competitive or the stabilizing role for inhibition in the circuit. These results generate testable predictions for perturbation experiments.

## Results

We consider circuits where two excitatory (E) populations integrate dedicated streams of sensory evidence to produce a categorical choice (Fig. 1a). In contrast to previous circuit models of decision-making with global inhibition, we include two inhibitory (I) populations which can inherit choice selectivity from excitatory neurons (Methods). The circuit dynamics are modeled using two-dimensional meanfield equations where the mean postsynaptic activation of the two excitatory (E_1_ and E_2_) populations are the dynamic variables^4^. The average strength of connections between the four choice selective populations is controlled by a specificity parameter Σ. For each of three connection classes (E to E, E to I, and I to E; Fig. 1b), Σ sets the balance of connection strengths between populations with the same and opposite choice selectivity (Fig. 1c), where (1 + Σ^*ij*^) is the strength of connections between populations selective for the same choice and (1 – Σ^*ij*^) is the strength of connections between populations selective for the opposite choice (for the presynaptic population i, either E or I, and the postsynaptic population j). We keep Σ^EE^ positive due to the importance of recurrent excitation in the function of these circuits^4^. Inhibitory choice selectivity is controlled by Σ^EI^ and is also positive because inhibitory neurons inherit choice and stimulus information from the excitatory neurons. Thus, inhibitory activity is not choice selective when Σ^EI^ = 0 because inhibitory neurons receive equal input from both excitatory populations. Inhibitory choice selectivity emerges as Σ^EI^ increases (Fig. 1d).

**Figure 1.**
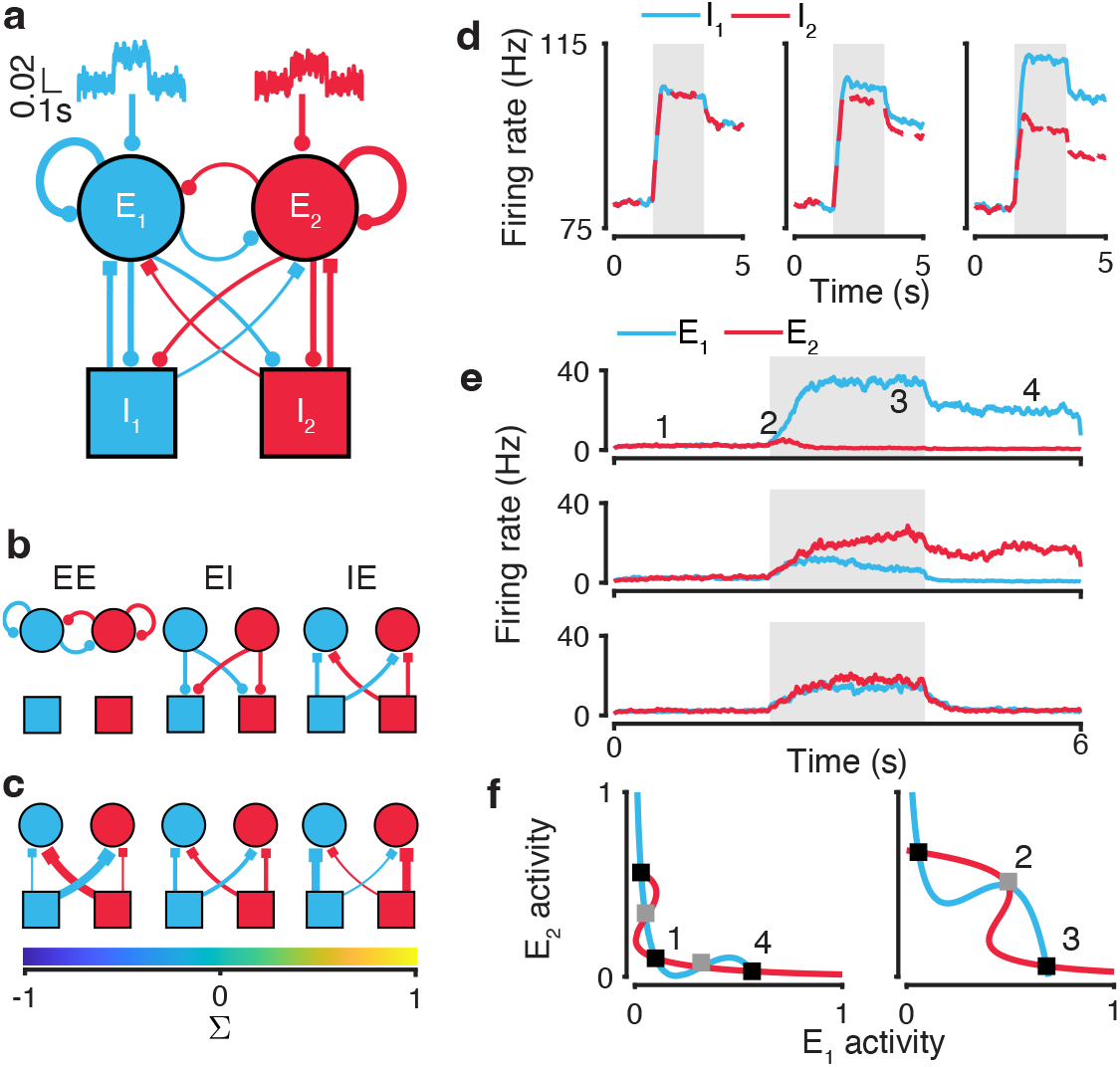
A mean-field circuit model of decision making with choice-selective inhibition. (**a**) The circuit diagram of the model with choice-selective excitatory and inhibitory populations. (**b**) The circuit model includes three connection classes: excitatory-excitatory (EE), excitatory-inhibitory (EI), and inhibitory-excitatory (IE). (**c**) The parameter Σ controls the specificity of connections between choice-selective populations. The output connections preferentially target neurons with the same choice preference when Σ is positive, and with the opposite choice preference when Σ is negative. (**d**) Σ^EI^ controls inhibitory choice selectivity. Firing rate of inhibitory populations for Σ^EI^ = 0 (left), Σ^EI^ = 0.05 (center), Σ^EI^ = 0.25 (right) are shown for an example trial with stimulus strength equal to 20. (**e**) Circuits report choices by elevating the firing rate of one excitatory population. Example trials showing E_1_ (blue) and E_2_ (red) population activity for stimulus strength equal to 20 (upper panel), and for stimulus strength equal to 0 on a completed (middle panel) and invalid trial (lower panel). Grey shading indicates stimulation period. Numbers indicate activity corresponding to fixed points in f. (**f**) Eight fixed points are required for decision-making dynamics in the circuit: five in the unstimulated phase-plane (left) and three in the stimulated phase-plane (right, stimulus strength is equal to 0). Lines show nullclines of E_1_ and E_2_ populations, black squares indicate fixed-point attractors, and gray squares indicate saddle-points for a circuit with nonselective inhibition.

For inhibitory choice selectivity to have any effect on circuit function, the outputs of inhibitory populations must be structured (i.e. Σ^IE^ ≠ 0; Fig. 1c). The specificity of inhibitory outputs Σ^IE^ can range between [−1, 1] with negative values favoring connections between E and I populations with opposite choice preference and positive values favoring connections between E and I populations with the same choice preference. Thus, the specificity of inhibitory output connectivity defines three circuit motifs: contraspecific for Σ^IE^ < 0, ipsispecific for Σ^IE^ > 0, and nonspecific for Σ^IE^ = 0.

In any decision-making circuit, inhibition concurrently fulfills two roles. The first is providing the substrate for competition between the excitatory populations, and the second is stabilizing the self-amplification driven by strongly recurrent excitatory populations. Both of these roles must be fulfilled for a circuit to function, but specific connections to and from inhibitory populations could enhance one of these roles (Fig. 1c). Specifically, ipsispecific inhibition can promote stabilizing feedback and contraspecific inhibition can maximize competition.

In response to an input stimulus, the circuit can produce different choice outcomes by changing the firing rates of the excitatory populations. Circuits report a choice by persistently raising the firing rate of one excitatory population at least 15 Hz above the other; trials where this separation does not occur are considered invalid (Fig. 1e). We also require that prior to the stimulus onset, the circuit maintains low, symmetric activation of excitatory neurons. These dynamics are governed by eight fixed points which are essential for the good decision-making behavior (Fig. 1e,f). Prior to stimulus onset, both excitatory populations maintain low symmetric activation, which is set by an attractor located near the origin of the unstimulated phase plane. Following stimulus onset, the firing rate for both populations increases as the system approaches a saddle point along the stable manifold which acts as a separatrix between two choice attractors. Following stimulus offset, the system returns to its unstimulated phase plane and the choice of the circuit is preserved by one of two working memory attractors.

### Inhibitory connection specificity expands the space of circuits that support decision making

Using the mean-field model, we investigated how the circuit’s ability to perform decision-making depends on the inhibitory connectivity structure. Specifically, we determined how choice-selective inhibition affects the presence of the eight fixed points governing decision-making behavior. We sampled the specificity parameter space to identify circuits which support these eight fixed points (Fig. 2a). We found that a broad range of circuit configurations can support decision making. There are two components of inhibitory choice selectivity which rely on specific connections to and from inhibitory populations. The first is the degree of choice selective firing by inhibitory neurons that is controlled by Σ^EI^. The second is the degree to which inhibitory populations have a specific effect on excitatory neurons that is controlled by Σ^IE^. We combine these two components into a selectivity index Σ^EI^Σ^IE^, which is negative for contraspecific and positive for ipsispecific circuits following the sign of Σ^IE^. The specificity of excitatory and inhibitory connections is highly correlated in circuits supporting decision making (Fig. 2b). When inhibition is nonselective (Σ^EI^ = 0) or nonspecific (Σ^IE^ = 0), the strength of recurrent excitation (Σ^EE^) is highly constrained and deviations from a narrow range leads to the loss of one of the essential fixed points (Fig. 2c). For circuits with selective inhibition, a wider range of Σ^EE^ will support decision making as long as a complementary inhibitory motif is present. For low Σ^EE^, the inhibitory motif must be contraspecific (Σ^EI^Σ^IE^ < 0, Fig. 2b and Fig. 2d left) and for high Σ^EE^ it must be ipsispecific (Σ^EI^Σ^IE^ > 0, Fig. 2b and Fig. 2d right). Contraspecific inhibitory motif can promote competition in circuits where excitatory feedback connections are insufficiently strong to amplify firing rate differences between choice selective populations. Ispispecific inhibitory motif can stabilize excitatory feedback to prevent inadvertent winner-take-all dynamics in the absence of stimulus in circuits with strong excitatory specificity. By enhancing either the competitive or stabilizing role, circuits with choice selective inhibitory populations can support decision making for a wider range of Σ^EE^ (Fig. 2b).

**Figure 2.**
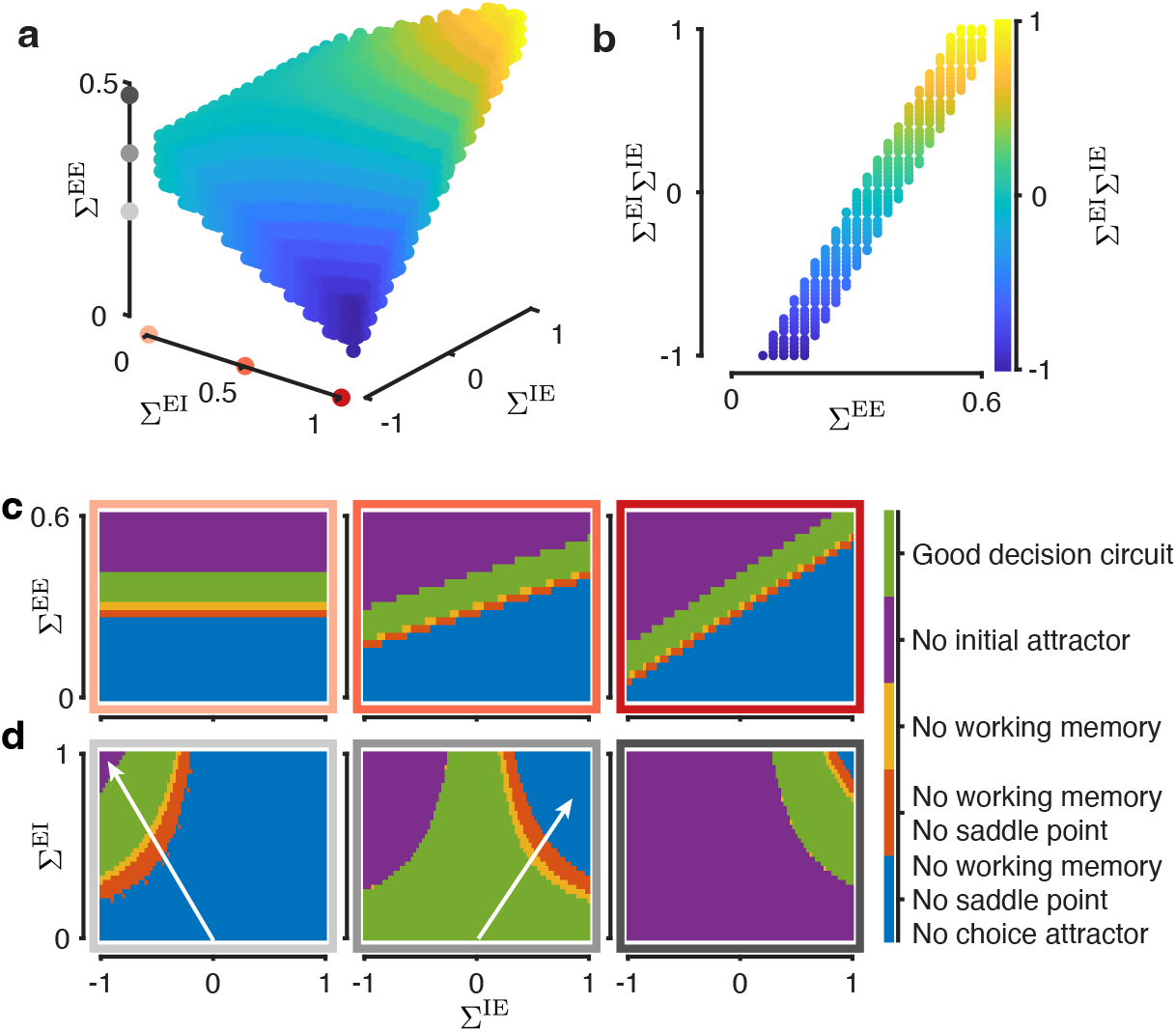
Choice-selective inhibition expands the space of circuits supporting decision making. (**a**) The volume in the selectivity parameter space where circuits have all fixed points necessary to support decisionmaking. Color indicates the inhibitory selectivity index Σ^EI^Σ^IE^. (**b**) For nonselective or nonspecific inhibition, circuits that support decision-making exist only for a narrow range of Σ^EE^. If inhibition is selective and specific, a broader range of Σ^EE^ becomes possible. Inhibitory specificity needs to be complementary to excitatory specificity, as reflected in the correlation between the inhibitory selectivity index and Σ^EE^ for circuits with all necessary fixed points. (**c**) With increasing inhibitory selectivity, a broader range of Σ^EE^ can support decision making, evident as a steeper slope of the region with all required fixed points (green, good decision circuit). Panels show slices through the parameter space in a at Σ^EI^ = 0.0 (left), Σ^EI^ = 0.5 (center), Σ^EI^ = 1.0 (right). Colored frames correspond to dots on the Σ^EI^ axis in a. (**d**) When Σ^EE^ is low inhibition must be contraspecific (left) and when Σ^EE^ is high inhibition must be ipsispecific (right). Slices through the parameter space in a at Σ^EE^ = 0.225 (left), Σ^EE^ = 0.35 (center), Σ^EE^ = 0.475 (right). Colored frames correspond to dots on the Σ^EE^ axis in a. Arrows indicate a sequential loss of fixed points described in the text.

The emphasis on competition or stability can also be seen in which fixed points are lost when connection specificity between excitatory and inhibitory populations are not complementary. When Σ^EE^ is low, nonspecific and ipsispecific circuits lack the fixed points representing choice both in the presence and absence of stimulation as well as the saddle point during the stimulus (Fig. 2d left, Fig. S1), because recurrent excitation is too weak to drive competition alone. Contraspecific inhibition paired with low Σ^EE^ restores these fixed points by emphasizing competition between populations selective for opposite choices. These fixed points emerge sequentially as the inhibitory motif becomes more contraspecific: first the choice attractors appear, followed by the saddle point, and finally by the working memory attractors (arrow in Fig. 2d left). For moderate Σ^EE^, nonspecific circuits have all eight necessary fixed points, but deviations to a contraspecific motif cause the loss of the attractor for the low initial state, whereas deviations to an ipsispecific motif cause the loss of the working memory attractors, then saddle point, and then choice attactors (arrow in Fig. 2d center, Fig. S1). For circuits with high Σ^EE^ to support decision making, inhibitory motif must be ipsispecific, as nonspecific and contraspecific circuits lack the initial low activation state attractor (Fig. 1d right, Fig. S1).

### Inhibitory motif controls the speed versus accuracy trade-off

The roles enhanced by contra- and ipsispecific inhibititory motifs lead to differences in performance of decision circuits. In circuits with moderate strengths of recurrent excitation, all three motifs can support decision making for the same Σ^EE^. We found that circuits with three inhibitory motifs differ in choice accuracy on difficult trials where stimulus strength is weak (Fig. 3a). Relative to a circuit with nonspecific inhibitory outputs (Σ^IE^ = 0), ipsipecific circuits are more accurate at classifying difficult stimuli but more often fail to separate the outputs sufficiently producing invalid trials (Fig. 3b). Contraspecific circuits, on the other hand, have lower accuracy for difficult stimuli. In addition, contraspecific circuits have a stimulus independent rate of trial failure attributable to trials where the firing rates of choice-selective populations separate prior to the stimulus onset (Fig. 3b), highlighting how these circuits are primed for competitive dynamics. It is well known that decision accuracy and reaction time are linked through the speed-accuracy trade-off, where longer integration times lead to more accurate decisions^22–24^. Ipsispecific circuits could be more accurate at the expense of speed, so we compared the average time it takes circuits to cross the decision threshold for each stimulus strength as a proxy for reaction time. Ipsispecific circuits do indeed arrive at choices more slowly than the less accurate contraspecific circuits (Fig. 3c). These differences in behavioral performance indicate a speed versus accuracy trade-off which is mediated by the specificity of connections between choice-selective populations in the circuit. These performance outcomes again highlight the roles enhanced by ipsispecific and contraspecific inhibition: the contraspeicific motif primes a circuit for competition, whereas the ipsispecific motif promotes stability lengthening integration times.

**Figure 3.**
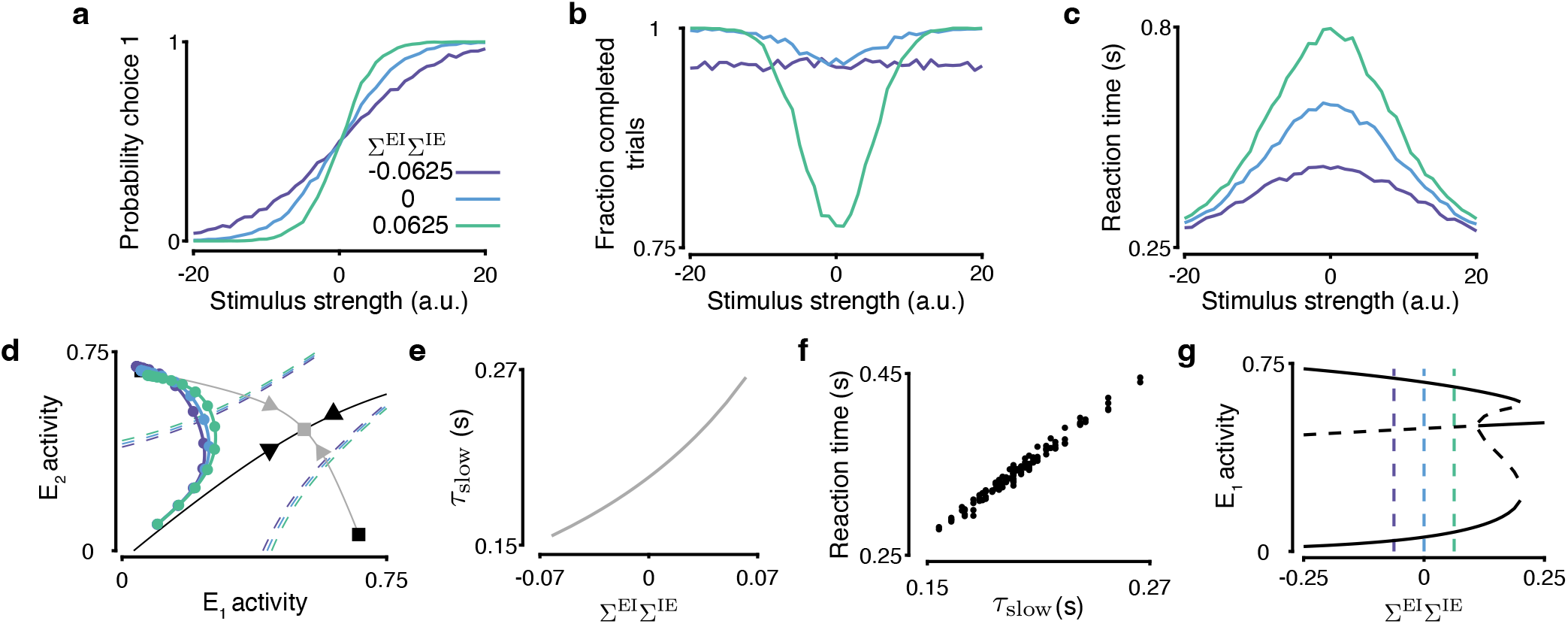
Inhibitory circuit motifs mediate the speed-accuracy trade-off in decision-making. (**a**-**c**) Contraspecific circuits are faster and less accurate, whereas ipsispecific circuits are slower and more accurate than nonspecific circuits. Psychometric functions (a), probability of trial completion (b), and chronometric functions (c) for circuits with different inhibitory motifs. (**d**) Contraspecific circuits deviate to a choice attractor earlier and faster than ipsispecific circuits. Single-trial trajectories are shown for three circuits with different inhibitory motifs, in 50 ms time steps (dots) for 0 stimulus strength. Noise is reduced by 50% for illustration clarity. The decision threshold for each circuit is shown by the dashed line. Squares indicate choice attractors, triangle indicates the saddle point. The stable (black) and unstable (red) manifolds of the saddle point are shown. (**e**) As the circuit motif changes from contra- to ipsispecific, the time constant of the unstable eigenvector of the saddle point *τ*_slow_ increases, indicating stabilization of dynamics and longer integration times. (**f**) The time constant *τ*_slow_ is tightly correlated with reaction time (shown for stimulus strength equal to 0). (**g**) The saddle point becomes an attractor for ipsispecific circuits with high Σ^EI^Σ^IE^. The bifurcation diagram for circuits driven by a stimulus of 0 strength shows the location of attractors (black solid line) and saddle points (black dashed line). Dashed vertical lines correspond to examples in a-d. In all panels Σ^EE^ = 0.32 and Σ^EI^ = 0.25.

We can understand the speed-accuracy trade-off between ipsi- and contraspecific circuits by analyzing the dynamics around the saddle point. Differences in these dynamics are seen by comparing single-trial trajectories of ipsi-, non-, and contrapecific circuits in response to the neutral stimulus (Fig. 3d). At the trial start, both choice-selective populations are symmetrically activated and the trajectory moves along the stable manifold toward the saddle point. Eventually, the circuit activity deviates to a choice attractor after approaching the saddle. Contra- and ipsispecific circuits differ in both how far along the stable manifold the activity progresses and how quickly it moves toward the choice attractor once it deviates. We can estimate how quickly the dynamics will leave the neighborhood of the saddle point with the time-constant *τ*_slow_, which is the time-constant of dynamics moving along the unstable manifold of the saddle point^4^. Changing the circuit motif from contraspecific to ipsispecific by increasing Σ^EI^Σ^IE^ leads to an increase in *τ*_slow_ (Fig. 3e) and slowing down the pace of decisions (Fig. 3f). The divergence of *τ*_slow_ indicates that ipsispecific inhibition stabilizes the saddle point until at high Σ^EI^Σ^IE^ a bifurcation occurs and the saddle point becomes an attractor with a symmetric high activity state (Fig. 3g). This bifurcation leads to the system stabilizing in a state where firing rates of two choice-selective populations do not sufficiently separate on neutral and difficult stimuli trials, a state where the circuit fails to produce a decision. Easy stimuli impose a stronger asymmetry on the phase plane^4^ allowing circuits with highly ipsispecific inhibition to make choices on easy trials (Fig. S2).

### Strong ipsispecific inhibition destabilizes working memory

Persistence of the decision after stimulus offset allows for a choice readout to be made even after a significant delay and is a hallmark of good decision-making in the circuit (Fig. 1d). Contraspecific and nonspecific circuits maintain a difference in excitatory firing rates of at least 15 Hz for a very long time following stimulus offset, whereas ipsispecific circuits exhibit a degradation of the choice readout (Fig. 4a). This behavior can be linked to the phase plane of the unstimulated circuit. Working memory is supported by two choice attractors that are separated by saddle points from the attractor with symmetric low activity state. The separation between the working memory attractors and the saddle points is smaller for more ipsispecific circuits (Fig. 4b). For highly ipsispecific circuits, working memory attractors are extinguished after merging with the saddle points (Fig. 4b).

**Figure 4.**
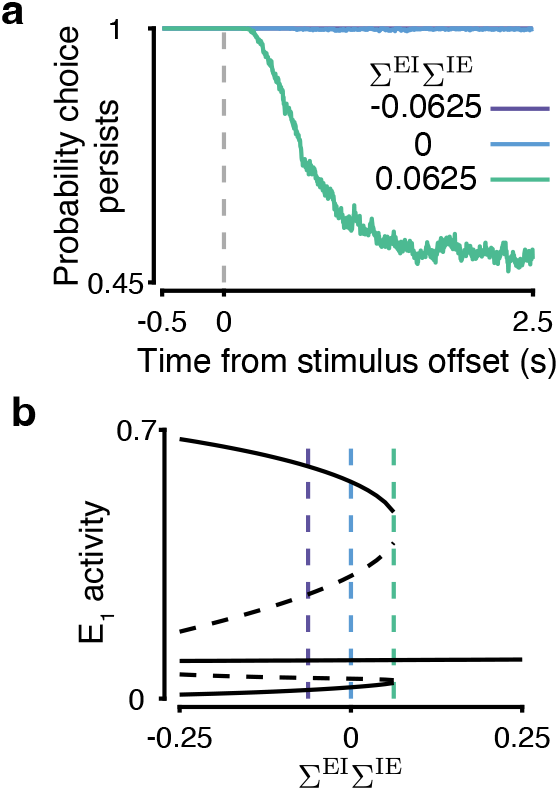
Strong ipsispecific inhibition destabilizes working memory attractors. (**a**) The probability of maintaining a choice after stimulus offset is diminished in ipsispecific circuits. (**b**) For ipsispecific circuits with high Σ^EI^Σ^IE^, working memory attractors are extinguished after merging with saddle points. The bifurcation diagram is for the same circuits as in Fig. 3g but in the absence of stimulus.

### Inhibitory choice selectivity in trained recurrent neural networks

So far, we used the mean-field approach to establish that choice-selective inhibition supports the function of decision-making circuits by enhancing a competitive or stabilizing role. Next, we wanted to test whether this result holds broadly by using another class of decision-making network models. We therefore trained excitatory-inhibitory recurrent neural networks (RNNs) to perform a decision-making task^25^ and then tested whether inhibitory choice-selectivity regularly emerges in these networks after training and whether the dependence between the excitatory and inhibitory specificity aligns with the two roles for inhibition. We used RNNs with 100 excitatory and 25 inhibitory units (Fig. 5a). Two input streams projected to all excitatory units through input weights. Two output variables were calculated as a weighted sum of excitatory unit activity. We trained RNNs to perform an identical decision-making task as the mean-field circuits by raising an output variable which corresponds to the input stream with a higher mean value. Networks were trained by back-propagation through time to minimize the mean squared error between the network outputs and predefined targets. For a given trial, a choice was recorded when the output variables became separated by a fixed threshold set to 0.25. Trials were considered invalid if the outputs separated prior to the stimulus, failed to maintain separation after stimulus offset, or separation was never achieved. We trained networks until the correct choice was made on 85% of trials in a 200 trial epoch (counting invalid trials as incorrect). One hundred and fifty networks reached this training threshold in 10, 4343 ± 9, 264 (mean ± s.t.d.) trials (Fig. 5b). Networks performed the task well, making errors and failing to complete trails only for difficult stimuli (Fig. 5c). Trained networks also took longer to make decisions when presented with a difficult stimulus, similarly to mean-field circuits (Fig. S3).

**Figure 5.**
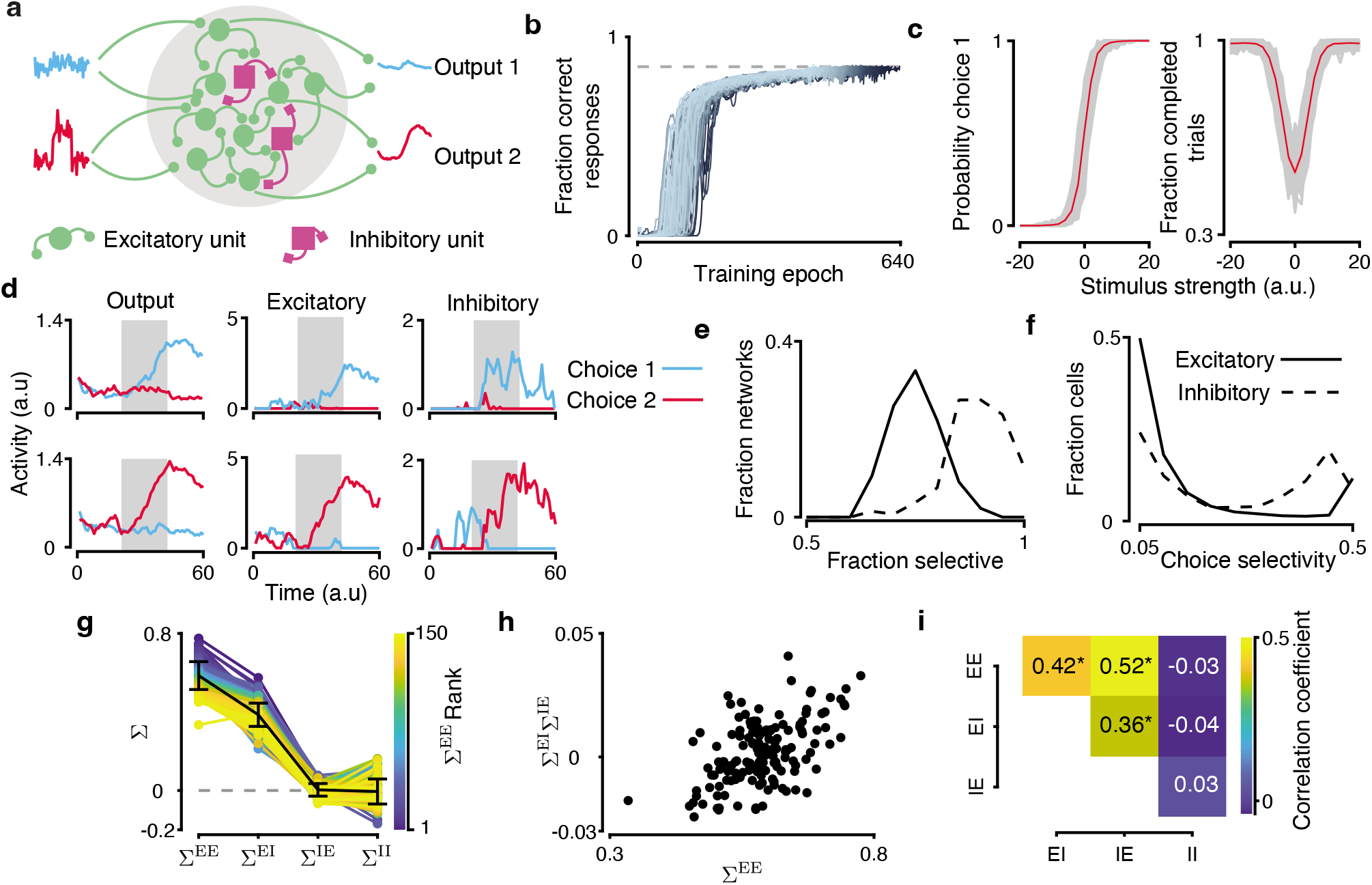
Inhibitory choice selectivity in trained recurrent neural networks. (**a**) RNNs are trained to compare two inputs and indicate which has higher mean by elevating the corresponding output. RNNs are composed of 100 excitatory and 25 inhibitory units. (**b**) We trained 150 RNNs to a consistent level of performance. RNN performance improved gradually during training. We stopped the training when the network performance reached 85% correct responses (grey dashed line). Lines show individual RNNs, color gradient indicates the network’s rank to reach 85% performance. (**c**) Psychometric functions (upper panel) and probability of trial completion (lower panel) for all trained RNNs (grey) and their average (red). (**d**) Excitatory and inhibitory units in trained RNNs display choice selectivity. Traces show the activity of the RNN outputs (left), two excitatory units (center), and two inhibitory units (right) on two example trials with choice 1 (upper row, stimulus strength −20) and choice 2 (lower row, stimulus strength 20). Grey shading indicates the stimulus period. (**e**) In trained RNNs, the overall choice selectivity is greater for inhibitory than excitatory units. Distributions show the choice-selectivity index across all units from all networks. (**f**) In trained RNNs, the fraction of units with significant choice selectivity is greater for inhibitory than excitatory units. Distributions show the fraction of selective units across networks. (**g**) Trained RNNs show a range of specificity parameters Σ for each connection class (colored lines - individual RNNs, black - mean ± s.t.d. across networks). Color indicates networks sorted by Σ^EE^. (**h**) In trained RNNs, the specificity index Σ^EI^Σ^IE^ is positively correlated with Σ^EE^. (**i**) In trained RNNs, excitatory-excitatory, excitatory-inhibitory, and inhibitory-excitatory specificity are correlated, whereas inhibitory-inhibitory specificity is uncorrelated with other connection classes. * indicates significant correlation.

We determined whether inhibitory neurons in these RNNs were choice selective. We classified recurrent units as choice selective using receiver operator characteristic (ROC) analysis (Methods)^21^. We constructed ROC curves by decoding network choice from a unit’s activity on the time-step following stimulus offset. To identify which units significantly modulated their firing rate to reflect choice, we compared the area under the ROC curve (AUC_ROC_) to a shuffle distribution generated from randomized trial labels (two-sided permutation test, *p* < 0.05). Units that were identified as choice selective increased activation following the onset of a stimulus corresponding to their preferred choice (Fig. 5d). Inhibitory units had overall higher choice selectivity than excitatory units, as measured by the selectivity index |AUC_ROC_ – 0.5| that can range from 0 to 0.5 (Fig. 5e, inhibitory 0.23 ± 0.17, excitatory 0.12 ± 0.16; mean ± s.t.d.; Wilcoxon rank-sum test *p* < 10^−10^). Also, the proportion of significantly selective units was higher for inhibitory than excitatory units (Fig. 5f, inhibitory 0.87 ± 0.07, excitatory 0.72 ± 0.06; mean ± s.t.d.; Wilcoxon Rank-Sum test *p* < 10^−10^). Thus, inhibitory unit activity contained overall more choice information than excitatory unit activity despite the fact that only excitatory units received stimulus input.

### Excitatory specificity aligns with ispi- and contraspecific inhibitory motifs in RNNs

Based on our mean-field model, we know that for choice-selective inhibition to impact circuit function, the connections from inhibitory to excitatory populations must be specific. Therefore, after identifying choice-selective units in RNNs, we sought to determine whether the connection specificity of excitatory-excitatory and excitatory-inhibitory pairs followed the relationship predicted by the mean-field model (Fig. 2b). To analyze the specificity of connections between choice-selective populations in the RNNs, we estimated the specificity parameter Σ from the weights of trained RNNs defined in the same way as for the meanfield model (Methods). Trained networks consistently had strong excitatory-excitatory (Σ^EE^ = 0.59 ± 0.07) and excitatory-inhibitory (Σ^EI^ = 0.39±0.06) specificity (Fig. 5g). This result is consistent with the constraint that inhibitory units inherit stimulus information from excitatory units to be choice or stimulus selective. Inhibitory-excitatory connections were nonspecific on average (Σ^IE^ = 3.6 × 10^−3^ ± 0.03) but their distribution showed both ipsispecific and contraspecific motifs. Unlike our mean-field model, specific connections could also emerge between inhibitory-inhibitory units during the training process in RNNs. Inhibitory-inhibitory connections were nonspecific on average with higher variation than inhibitory-excitatory connections (Σ^π^ = −5.0 × 10 – 3 ± 0.06). Confirming the trend predicted by the mean-field model, excitatory specificity Σ^EE^ was correlated with the inhibitory specificity index Σ^EI^Σ^*IE*^, where networks with stronger recurrent excitation were ipsispecific and networks with weaker recurrent excitation were contraspecific (Pearson’s *r* = 0.53, *p* < 10^−10^; Fig. 5h). When comparing the connection classes individually, we found positive correlations between excitatory-excitatory, excitatory-inhibitory, and inhibitory-excitatory specificity (Fig. 5i). Inhibitory-inhibitory connection specificity was not significantly correlated with any other connection class. The higher variance and negligible correlation with other connection classes suggest that the specificity of inhibitory-inhibitory connections was unconstrained in these networks. These results show that RNNs utilize choice selective inhibition to compensate for variation in excitatory-excitatory specificity.

To further test the relationship between the excitatory and inhibitory specificity, we trained additional sets of RNNs with higher or lower excitability of excitatory units. In the mean-field model, lower (higher) excitatory gain can be compensated by either an increase (decrease) in excitatory connection specificity or by strengthening of the contraspecific (ipsispecific) motif. Accordingly, we expect that changing the activation function slope of the excitatory units in RNNs should either shift the excitatory-excitatory specificity against the direction of the gain change or shift the inhibitory specificity towards contraselective (for lower slope) or ipsielective motif (for higher slope). We trained two additional sets of networks with hypoexcitable (slope 0.5) or hyperexcitable (slope 1.5) excitatory units. Changing the excitability of excitatory units led to large shifts in Σ^EE^ without changing the distribution of inhibitory specificity (Fig. S4). In these networks, Σ^EE^ and Σ^EI^Σ^IE^ were still correlated, with higher Σ^EE^ leading to higher Σ^EI^Σ^IE^. These results indicate that excitatory-excitatory specificity is a higher leverage parameter that RNNs use as the most effective path to compensate for changes in the excitability of excitatory units. This observation is consistent with the effect of changes in Σ^EE^ on the dynamics in the mean-field model. For both reaction-time and *τ*_slow_, changes in Σ^EE^ are far more effective than changes in inhibitory specificity (Fig. S6). In both the mean-field and RNN models, excitatory-excitatory specificity has a larger effect than inhibitory specificity and is the main lever circuits use to compensate for changes in neural parameters.

### Perturbing inhibitory neuron activity highlights two roles for inhibition in decision making

Using the mean-filed and RNN models, we established how contra- and ipsispecific inhibitory motifs enhance two different roles for inhibition in decision making circuits. To further probe these roles, we next considered how circuits respond to perturbations of inhibitory neuron activity. We used perturbations that equally targeted all inhibitory neurons irrespective of their choice selectivity by driving them with a nonspecific input Δ*ν*_0,I_ (Fig. 6a). Such perturbations could be realized in optogenetic experiments. In circuits where the competitive role of inhibition dominates, we expect that enhancing inhibitory activity should speed up dynamics whereas suppressing inhibition should slow them down (Fig. 6b). Vice versa, in circuits where the stabilizing role of inhibition dominates, we expect that enhancing inhibitory activity should slow dynamics down and suppressing inhibition should speed them up (Fig. 6b). Because *τ*_slow_ provides a readily available estimate of the pace of dynamics in the mean-field model, we calculated *τ*_slow_ for varying nonspecific input to inhibitory neurons *ν*_0,I_. We found that depending on the baseline level of inhibitiory activity both regimes are possible in the mean-field circuit: one where competitive role dominates and one where stabilizing role dominates (Fig. 6c). Around a low baseline value of inhibitory activity (*ν*_0,I_ = 11.5 in Fig. 6c), contra-, ipsi-, and nonspecific circuits respond to perturbations similarly, such that enhancing inhibition (Δ*ν*_0,I_ > 0) leads to a decrease in *τ*_slow_, i.e. faster dynamics. Around a high baseline value of inhibitory activity (*ν*_0,I_ = 14 in Fig. 6c), all circuits respond in the opposite way, such that enhancing inhibition increases *τ*_slow_. These two regimes–a low inhibition and a high inhibition regime–differ in which role of inhibition dominates: competitive or stabilizing, respectively. The inhibitory motif (contra-, non-, or ipsispecific) further shifts this emphasis within the constraints of each regime. These regimes can be identified via perturbations by characterizing how the circuit dynamics respond to changes in inhibitory tone.

**Figure 6.**
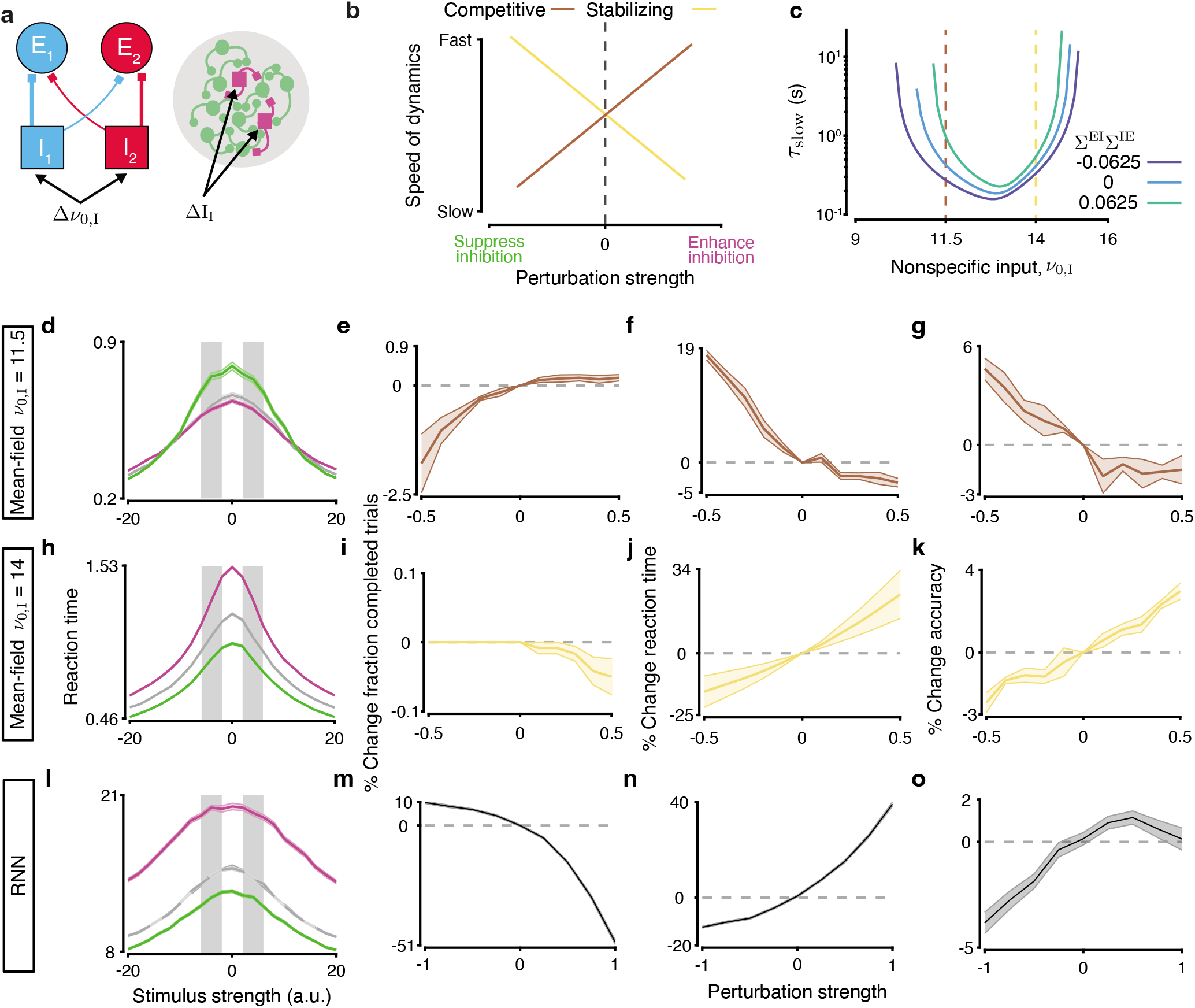
Perturbations to inhibitory activity reveal regimes where stabilizing and competitive inhibition dominate. (**a**) We perturbed mean-field models by delivering a nonspecific input Δ*ν*_0,I_ to all inhibitory neurons during stimulus period. Perturbations were similarly delivered to RNNs by adding a constant input ΔI_I_ to all inhibitory units during the stimulus. (**b**) Circuits where competitive (brown) or stabilizing (yellow) inhibition dominates are predicted to have diverging responses to perturbations in inhibitory activity. (**c**) Dependence of *τ*_slow_ on the baseline input to inhibition *ν*_0,I_ reveals two inhibitory regimes: a low inhibition regime where competitive inhibition dominates and a high inhibition regime where stabilizing inhibition dominates. Contra- or ipsispecific inhibitory motifs shift the emphasis within these regimes (e.g., *τ*_slow_ is always longer for ipsi-than contraspecific circuits). (**d**-**k**) The effects of inhibitory perturbations in the meanfield model differ depending on the baseline value *ν*_0,I_. (**d**-**g**) Around a low baseline (*ν*_0,I_ = 11.5, brown line in c), enhancing inhibition speeds up reaction times (magenta line in d; f), increases the rate of trial completion (e), and decreases accuracy (g), whereas suppressing inhibition produces the opposite effects, e.g., slows down reaction times (green line in d; j). Results are shown for nonspecific circuits. Gray areas in **d** indicate stimulus strengths used to calculated the values in **e**-**g**. (**h**-**k**) Same as d-g for the high inhibition regime (*ν*_0,I_ = 14, yellow line in c). Perturbations of inhibitory activity produce the reversed effects. (**l**-**o**) Same as h-k for perturbations of inhibitory neurons in RNNs. RNN’s response to perturbations mirrors the effects in the mean-field model in the stabilizing regime (c.f. h-k). Enhancing inhibition in RNNs slows down reaction times, decreases the rate of trial completion, and increases accuracy.

To confirm the existence of competitive and stabilizing regimes, we perturbed the mean-field circuits around the low and high baseline values of the inhibitory activity. We enhanced or suppressed inhibition during the stimulus period of a trial and measured changes in the circuit performance. We constructed a set of metrics to quantify changes in the fraction of completed trials, reaction time, and choice accuracy relative to the unperturbed circuit for all stimulus strength. The effects of these perturbations followed the predictions from the calculation of *τ*_slow_ (Fig. 6d-k). Enhancing inhibition decreased reaction time in the low inhibition regime, but increased reaction time in the high inhibition regime (cf. Fig. 6d,f and h,j). Consistent with the slowing effects of the perturbation, circuits in the high inhibition regime failed more often to complete trials (Fig. 6e) and became more accurate (Fig. 6g) when inhibition was enhanced. Circuits in the low inhibition regime showed the opposite behavior (Fig. 6h-k). Thus, by perturbing inhibitory neuron activity we can determine whether the competitive or stabilizing inhibition dominates in a circuit.

We then delivered enhancing or suppressing perturbations to inhibitory units in trained RNNs during the stimulus period to identify in which inhibitory regime these networks operate. Enhancing inhibition increased reaction times, reduced the fraction of completed trials, and increased accuracy, consistent with these RNNs operating in the stabilizing inhibition regime (cf. Fig. 6h-k and l-o).

## Discussion

We showed that choice selectivity of inhibitory neurons can affect the function of decision making circuits by enhancing one of two roles for inhibition: facilitating competition or stabilizing recurrent excitation. In the mean-field model, choice selective inhibition and specific connections from inhibitory to excitatory populations broaden the parameter space of circuits that support decision-making. For the range of excitatory connection specificities supporting both ipsispecific and contraspecific inhibitory circuits, the speed and accuracy of decisions tightly depend on whether the ipsi- or contraspecific inhibitory motif is present. Inhibitory choice selectivity also emerges in RNNs trained to perform a decision-making task, and the specificity of excitatory and inhibitory connections within trained RNNs is correlated, consistent with the mean-field model predictions. Perturbations suppressing or enhancing all inhibitory neurons reveal the existence of two regimes in the mean-field model: (i) a low-inhibition regime where the competitive role dominates, and (ii) a high-inhibition regime where stabilizing role dominates. In trained RNNs, perturbations of all inhibitory neurons indicate that these networks operate in the stabilizing inhibition regime.

### Selective inhibition broadens the range of circuits capable of decision-making

Decision-making circuits with non-selective inhibition exist only within a narrow range of excitatory-excitatory connection specificity. When inhibitory neurons inherit choice-selectivity from excitatory neurons and also project to excitatory neurons via specific connections, a broad range of circuit configurations can support decision-making. In circuits capable of decision-making, the correlation between the specificity of excitatory (Σ^EE^) and inhibitory connections (Σ^EI^Σ^IE^) reveals how the contra- and ipsispecific motifs enhance one of two roles for inhibition: facilitate competition between populations coding for opposite choices or stabilize amplification driven by strongly recurrent excitation. When Σ*^EE^* is low and excitatory populations alone cannot drive selective activation, contraspecific inhibitory motifs support decision-making by maximizing competition. Conversely, when Σ^*EE*^ is high and excitatory self amplification becomes unstable, ipsispecific inhibitory motifs stabilize firing rates.

The categorical output of decision-making circuits is thought to be driven by strongly selective excitatory to excitatory selectivity with the evidence accumulation based on amplification through N-Methyl-D-Aspartate receptors^2, 4^. In these models the specificity of excitatory connections is sufficient to drive competition and selective activation. We found that deviations from a narrow range of Σ^EE^ require complementary inhibitory circuitry. When recurrent excitatory specificity is low, contraspecific inhibition is required to form the attractors needed for decision-making computation. This mechanism was described in circuits where excitatory populations have limited capacity for amplification, such as the midbrain circuit in the owl^26^. On the other hand, when recurrent excitatory specificity is high, the strong excitatory feedback amplification needs matching ipsispecific inhibition to stabilize the circuit. This mode of inhibitory selectivity is known to improve stability and robustness of a circuit to perturbations^17, 27^.

We found a similar relationship between excitatory and inhibitory connection specificity in RNNs suggesting the balance between competitive and stabilizing inhibition is a general principle in E-I networks. While specific connections between excitatory and inhibitory units were clearly important for the decision-making function in our networks, connections between inhibitory units appeared unconstrained. RNNs are increasingly often used to develop theories of how neural circuits perform computations^25, 28, 29^. Some studies trained RNNs under the constraint that units have either exclusively excitatory or exclusively inhibitory outputs (Dale’s law)^25, 30^. Studies of E-I RNNs which focus on the impact of inhibitory connections show that specificity of inhibitory-inhibitory connections can be critical to circuit function^29^. The apparent difference in the importance of inhibitory-inhibitory selectivity between our networks and previous work could result from differences in the training procedures^31^. We observed a large impact of RNN training hyperparameters on the emerging circuit structure. Future work is needed to understand how details of training influence the emerging circuit structure and computations performed by RNNs.

### Selective inhibition may be a general feature of neural circuits

Our results show that selective inhibition can have a marked effect on the function of neural circuits. Many models of categorical decision-making rely on a nonspecific pool of inhibitory neurons to enforce winner-take-all competition between excitatory neurons^2, 3^. While these models reproduce the dynamics of decision-making circuits they do not fully account for the diversity of interneurons within the cortex. Cortical inhibitory neurons show selective activation in many modalities including primary sensory^13, 17, 32, 33^ and association areas^19–21^. Moreover, choice-selectivity of parietal inhibitory neurons is equal to that of excitatory neurons during an audio-visual discrimination task^21^.

In the mean-field model, we assume that choice selectivity of inhibitory neurons arises from specific connections from choice-selective excitatory neurons (Σ^EI^ in our model). While it is possible that choice selectivity could arise from external inputs to interneurons^34^ or even from random connections between excitatory and inhibitory neurons^35^, most circuit models assume stimulus information is exclusively provided by inputs to excitatory neurons. Inhibitory choice-selectivity also emerged in our RNNs trained to perform 2AFC task^25^. In our RNNs, inhibitory units can only inherit stimulus or choice information through specific connections from excitatory populations, unlike in other trained RNNs^29^.

### Impact of inhibitory circuitry on decision-making performance

The core computation of the model is the selective activation of a single excitatory population when the stimulus is presented and a mechanism to integrate stimulus information before diverting to a choice attractor. By enhancing stability, ipsispecific circuits lengthen the period when a circuit can maintain mutual activation of populations encoding competing choices, thus increasing the integration window which leads to more accurate stimulus classifications. Contraspecific circuits, primed for competition, minimize the integration period which increases error frequency.

In attractor networks, modulation of *τ*_slow_ for controlling the speed and accuracy of decisions can arise from other mechanisms than inhibitory output specificity. In the model with nonspecific inhibition, *τ*_slow_ increases with stimulus difficulty^4^ and can be also modulated via top-down excitation^36^. A key difference between controlling *τ*_slow_ via inhibitory motif versus top-down excitation is that the location of the saddle point is unaffected by Σ^IE^ whereas increasing top-down excitation shifts the saddle towards the origin, effectively acting as a collapsing decision-bound^36^. Top-down excitation can be adjusted rapidly from one trial to the next to match the decision’s speed and accuracy to the task demands. Could the inhibitory motif also be dynamically changed to meet changing task requirements? Modulation of the speed-accuracy trade-off through changes of the inhibitory motif may be mediated by activation or inactivation of inhibitory subpopulations connected in either a contraspecific or ipsispecific pattern (representing a shift in Σ^IE^ for the circuit as a whole).

Selective neuromodulatory control of genetically identifiable inhibitory subtypes may provide for control of inhibitory motifs. Inhibitory subtypes have distinct connectivity patterns to neighboring excitatory neurons: fast-spiking cells have far more reciprocal connections to excitatory neurons than adapting interneurons^16^. A shift in output specificity could be mediated through top-down activation of inhibitory subnetworks or through neuromodulation of distinct inhibitory subtypes such as PV^+^, SOM^+^, or VIP^+^. Acetylcholine has layer-dependent effects on the responsiveness of both regular spiking and fast spiking neurons in the visual cortex, which could differentially activate distinct inhibitory motifs on behaviorally relevant timescales^37–39^. Additionally, acetylcholine can reduce the release of inhibitory neurotransmitters in cortical neurons^40^, thus directly affecting inhibitory connectivity.

### Two roles for inhibition in decision-making

We show that choice selective inhibition can support one of two known roles for inhibition in decision-making circuits: facilitating competition or stabilizing excitatory feedback. Both these roles are simultaneously fulfilled by inhibition in any decision making circuit. By enhancing one of these roles, different inhibitory motifs expand the range of excitatory specificity which can support decision-making. The impact of circuit motif on the speed-accuracy trade-off reinforces this idea, as contraspecific inhibition promotes competition making decisions faster, whereas ipsispecific inhibition promotes stability slowing down decisions. Enhancing activity of all inhibitory neurons can shift the circuit from a regime where the competitive role dominates to a regime where the stabilizing role dominates regardless of which inhibitory motif is present. This echos results which find shifts in E/I balance can induce leaky or unstable integration^41^. The stabilizing and competitive regimes can be differentiated by the behavioral response to perturbations of inhibitory activity. Perturbations during reaction time tasks should reveal which inhibitory role is dominant *in vivo*. The balance of these two roles is critical for circuits to perform decision tasks, and shifts in this balance could align dynamics with changing task requirements. More experimental work is needed to uncover how inhibitory subnetworks strike this balance in the cortex. Specifically, whether functional selectivity is constrained to certain inhibitory subtypes and whether inhibitory neurons are recruited to perform a task in a state dependent manner are important questions for future work.

## Methods

### Mean-field model

We model the mean-field activity of the circuit through two variables representing the activations of N-methyl-D-aspartate (NMDA) conductances (in terms of fraction of channels open) onto two choice-selective excitatory populations^4^. The dynamics of the activation variable for population *i* (*i* ∈ {1, 2}) are governed by:

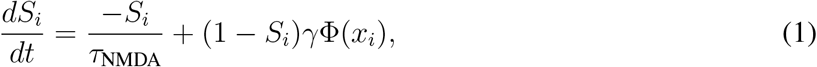

where *τ*_NMDA_ = 0.1 s and *γ* = 0.641. The non-linear function Φ transforms input current *x_i_* [nA] into firing rate:

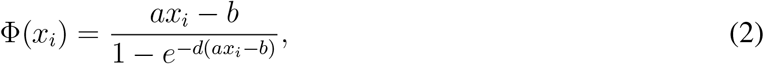

where *a* = 270 nC^−1^, *b* = 108 Hz, and *d* = 0.154 s. The input to population *i* is:

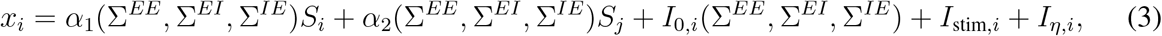

where index *j* refers to the other excitatory population. The complexity of the circuit structure, including interactions between all selective and nonselective excitatory and inhibitory neurons, is collapsed into two-dimensional model through the variables *α*_1_, *α*_2_, *I*_0, *i*_ as described in the section Circuit structure below.

The stimulus *I*_stim, *i*_ is defined as an increase in the rate of external excitatory inputs to choice-selective excitatory neurons of magnitude *μ*. We define the strength of evidence for one versus the other choice as stimulus coherence *c*, which can range between −100% and 100%. For population *i* the stimulus is then defined as:

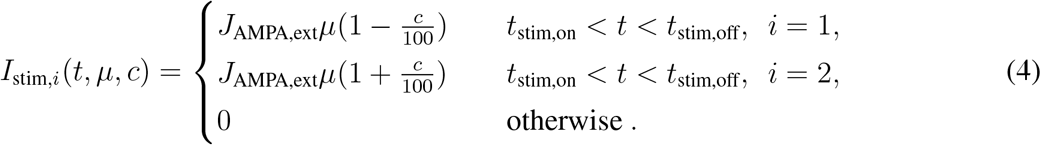

For all cases, we set *μ* to 40 Hz. Noise is introduced through the inputs *I_η,i_* to the two excitatory populations filtered through fast synaptic activation of *α*-amino-3-hydroxy-5-methyl-4-isoxazolepropionic acid (AMPA) receptors:

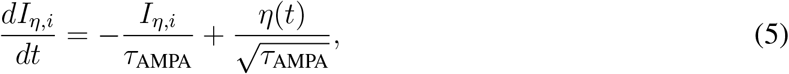

where *τ*_AMPA_ is 0.002 s and *η*(*t*) is a white Gaussian noise with zero mean and standard deviation 0.02 nA. We performed numerical simulations using the Euler method with a 2 ms time step.

### Circuit structure

We derived two-dimensional mean-field equations, which model the dynamics of the entire circuit through the effective interaction strengths *α*_1_, *α*_2_ between the two excitatory populations, and the background currents *I_0, *i*_*. This reduced model is based on approximating the firing rates of all three inhibitory populations (two choice-selective and one nonselective) and of the nonselective excitatory population as linear functions of their inputs. Thus, the firing rates of these populations change linearly in response to changes in the firing rates of the two explicitly modeled excitatory populations E_1_ and E_2_ ^4^. We define *α*_1_ as a term which describes how activity *S*_1(2)_ from the excitatory population E_1(2)_ filters through the circuit (i.e. via E_2(1)_, E_0_, I_0_, I_1_, I_2_, and feeding back onto itself) to impact its own firing rate. Similarly, *α*_2_ describes how the activity *S*_1(2)_ filters through the circuit to impact the firing rate of the opposite excitatory population. I_0, *i*_ describes the net input from the population activity that does not depend on the activity of E_1_ or E_2_. Thus, this model accounts for interactions between all six populations with only two dynamical system equations Eq. (1).

We parametrized connection specificity between choice-selective populations by Σ*^JK^* between presynaptic population *J* and postsynaptic population *K*. The index *J,K* ∈ {E,I} defines neuron type as excitatory or inhibitory. We translate Σ*^JK^* to a synaptic weight under a constraint that the total input to each population remains constant for all values of Σ*^JK^*. To this end, we defined an intermediate weight 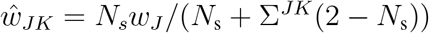, where *N_s_* = 2 is the number of competing choice-selective populations and *w_E_* = *w_I_* =1. We then set connection weights between populations with the same choice selectivity to 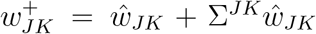 and between populations with opposite selectivity to 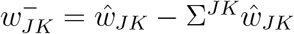. We can rewrite Σ in terms of *w*^+^ and *w*^−^ as:

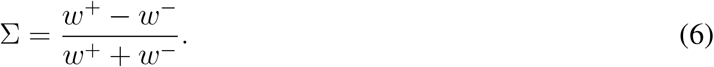

Connections to and from non-selective neurons were held at *w_J_* = 1. This definition enforces that all neurons receive the same total input weight for any value of Σ*^JK^*. We set the selectivity parameter ∑^EE^ = 0.32 as in Ref.^2, 4^, except in Figs. 1, 2. We set Σ*^EI^* = 0.25 except in Figs. 1, 2.

The effective interaction strengths *α*_1_ describes the recurrent feedback from an excitatory population’s activity onto itself fed through other populations in the circuit. This term consists of four components *α*_1_ = λ_1_(*α*_1*a*_ + *α*_1*b*_ + *α*_1*c*_ + *α*_1*d*_):

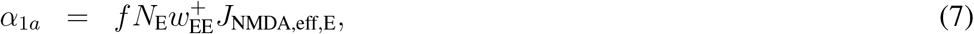

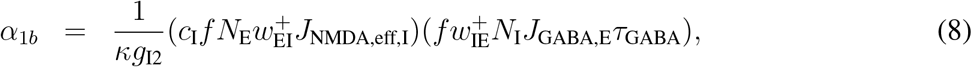

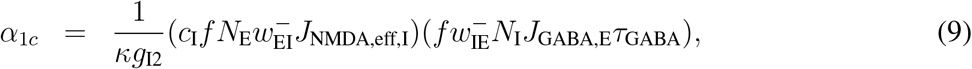

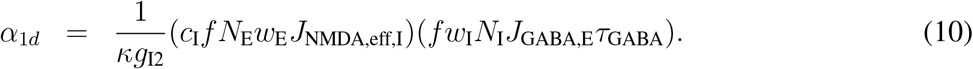

These components of *α*_1_ account for the effect of an excitatory population’s activity on its own activity filtered via (a) direct self-coupling, (b) the activity of the inhibitory population with the same choice selectivity, (c) the activity of the inhibitory population with the opposite choice selectivity, and (d) the activity of nonselective inhibitory neurons. Similarly, *α_2_* describes the influence of one excitatory population’s activity onto the other fed through all other populations in the circuit and also consists of four components *α_2_* = λ_2_(*α*_2*a*_ + *α*_2*b*_ + *α*_2*c*_ + *α*_2*d*_):

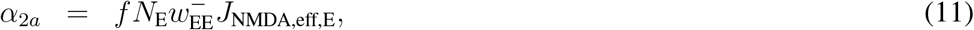

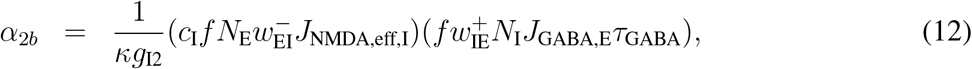

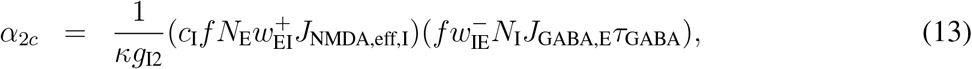

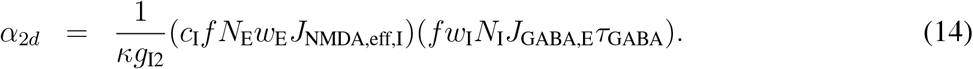

The components of *α*_2_ account for the effect on an excitatory population’s activity from the oppositely selective excitatory population’s activity filtered via (a) direct coupling, (b) the activity of the inhibitory population with the same selectivity, (c) the activity of the inhibitory population with the opposite selectivity, and (d) the activity of nonselective inhibitory neurons. The effects of nonselective neurons and external background inputs are described by *I*_0, *i*_ = λ*_I_*(*I*_0, *ia*_ + *I*_0, *ib*_ + *I*_0, *ic*_ + *I*_0, *id*_):

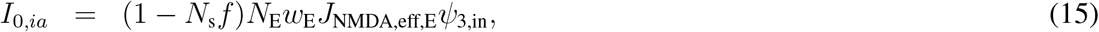

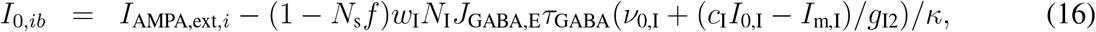

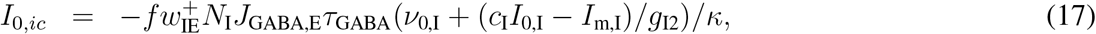

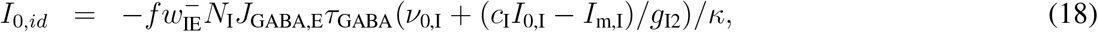

where:

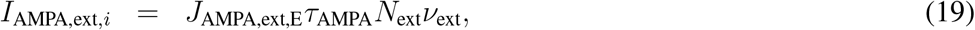

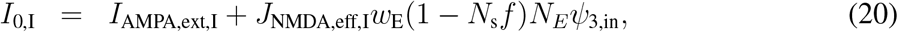

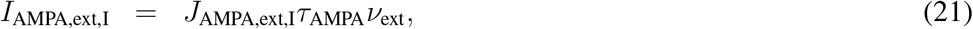

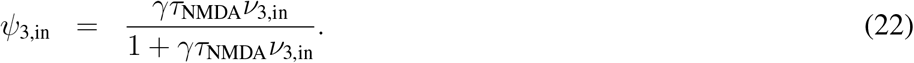

These terms account for the input to the excitatory population E*_i_* from the nonselective excitatory population filtered via (a) direct coupling, (b) the nonselective inhibitory population, (c) the inhibitory population with the same choice selectivity, (d) the inhibitory population with the opposite selectivity. The term *ψ* accounts for the NMDA activation of nonselective excitatory neurons. We calculated the firing rate of inhibitory populations as Φ_I,1(2)_ = *α*_1,I_*S*_1(2)_ + *α*_2,I_*S*_2(1)_ + *I*_0,II_, where:

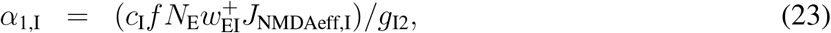

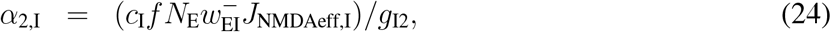

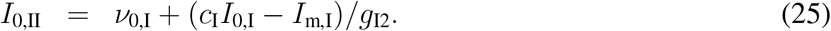

All parameter values are provided in Table 1.

**Table 1.** Mean-field model parameters.

| Parameter | Value | Description |
| --- | --- | --- |
| Mean-field physiological constants |  |  |
| $N_E$ | 1600 | Number of excitatory neurons |
| $N_I$ | 400 | Number of inhibitory neurons |
| $N_{\text{ext}}$ | 800 | Number of external inputs |
| $N_s$ | 2 | Number of possible choices/ choice selective populations |
| $\tau_{\text{NMDA}}$ (s) | 0.1 | Slow excitatory synaptic time constant |
| $\tau_{\text{AMPA}}$ (s) | 0.002 | Fast excitatory synaptic time constant |
| $\tau_{\text{GABA}}$ (s) | 0.005 | Inhibitory synaptic time constant |
| $\gamma$ | 0.641 | Firing-rate to NMDA activation scaling factor |
| $I_{\text{mI}}$ (Hz) | 177 | Inhibitory f-I curve intercept |
| $c_I$ (Hz/nA) | 615 | Inhibitory f-I curve slope |
| $g_{I2}$ | 2 | Inhibitory f-I curve scaling factor |
| $\nu_{0,I}$ (Hz) | $\in [9, 16]$ | Rate of background input to inhibitory neurons |
| $\nu_{\text{ext}}$ (Hz) | 3 | Rate of background input to selective excitatory neurons |
| $\nu_{3,\text{in}}$ (Hz) | 2 | Rate of background input to non-selective excitatory neurons |
| $V_E$ (mV) | -53.4 | Excitatory neuron resting potential |
| $V_I$ (mV) | -52.1 | Inhibitory neuron resting potential |
| $E_E$ (mV) | 0.0 | Excitatory synapse reversal potential |
| $E_I$ (mV) | -70.0 | Inhibitory synapse reversal potential |
| $g_{E,\text{rec,NMDA}}$ ($\mu\text{S}$) | $1.95 \times 10^{-4}$ | Maximum recurrent NMDA conductance, excitatory neurons |
| $g_{I,\text{rec,NMDA}}$ ($\mu\text{S}$) | $1.02 \times 10^{-4}$ | Maximum recurrent NMDA conductance, inhibitory neurons |
| $g_{E,\text{rec,GABA}}$ ($\mu\text{S}$) | 0.130 | Maximum recurrent GABA conductance, excitatory neurons |
| $g_{I,\text{rec,GABA}}$ ($\mu\text{S}$) | 0.0084 | Maximum recurrent GABA conductance, inhibitory neurons |
| $g_{E,\text{ext,AMPA}}$ ($\mu\text{S}$) | $2.1 \times 10^{-3}$ | Maximum external AMPA conductance, excitatory neurons |
| $g_{I,\text{ext,AMPA}}$ ($\mu\text{S}$) | $1.62 \times 10^{-3}$ | Maximum external AMPA conductance, inhibitory neurons |
| $J_{\text{AMPA,ext}}$ (nA/Hz) | $5.2 \times 10^{-4}$ | External stimulus current due to a single input event |
| $\lambda_1$ | 1.6719 | Scaling factor for $\alpha_1$ |
| $\lambda_2$ | 1.8844 | Scaling factor for $\alpha_2$ |
| $\lambda_I$ | 0.9229 | Scaling factor for $I_{0,i}$ |
| Mean-field derived constants |  |  |
| $\kappa$ | $1 + \frac{c_I}{g_{I2}} N_I J_{\text{GABA,I}} \tau_{\text{GABA}}$ | Linearized factor for inhibitory neurons |
| $J_{\text{GABA,E}}$ (nA) | $-g_{E,\text{rec,GABA}}(E_I - V_E)$ | Effective GABA current, excitatory neurons |
| $J_{\text{GABA,I}}$ (nA) | $-g_{I,\text{rec,GABA}}(E_I - V_I)$ | Effective GABA current, inhibitory neurons |
| $J_{\text{AMPA,ext,E}}$ (nA) | $g_{E,\text{ext,AMPA}}(E_E - V_E)$ | Effective AMPA current, excitatory neurons |
| $J_{\text{AMPA,ext,I}}$ (nA) | $g_{I,\text{ext,AMPA}}(E_E - V_I)$ | Effective AMPA current, inhibitory neurons |
| $J_{\text{NMDAeff,E}}$ (nA) | $\frac{g_{E,\text{rec,NMDA}}(E_E - V_E)}{1 + \frac{1}{3.57} e^{-0.062V_E}}$ | Effective NMDA current, excitatory neurons |
| $J_{\text{NMDAeff,I}}$ (nA) | $\frac{g_{I,\text{rec,NMDA}}(E_E - V_I)}{1 + \frac{1}{3.57} e^{-0.062V_I}}$ | Effective NMDA current, inhibitory neurons |

### Phase plane and bifurcation analysis

We analyzed the mean-field model to find null-clines and fixed points using MatLab’s fsolve function with the Levenberg-Marquant algorithm and a tolerance of 1 × 10^−6^. To identify the stability of the fixed points, we computed the Jacobian matrix analytically and found its eigenvalues numerically using the eig() function in MatLab. For the saddle points, *τ_slow_* is the positive eigenvalue of the Jacobian matrix.

### Recurrent neural network models

Recurrent neural networks (RNNs) were composed of 100 excitatory and 25 inhibitory units. The dynamics of these networks were governed by the equations:

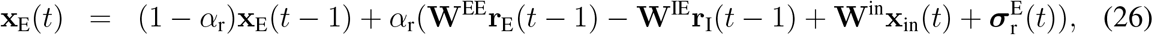

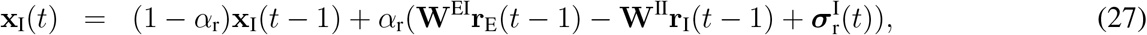

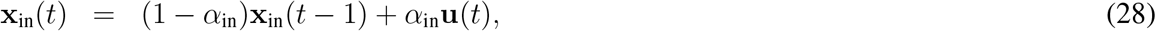

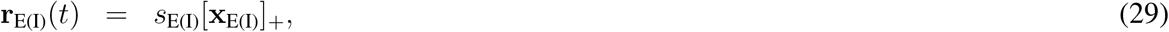

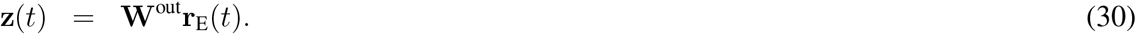

Here **x**_E_ and **x**_I_ are the vectors of activation variables for excitatory and inhibitory units, respectively. **r**_E_ and **r**_I_ are the corresponding activities after applying the rectified linear (RELU) nonlinearity 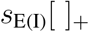, where *s*_E(I)_ sets the excitability of the excitatory or inhibitory units. **x**_in_ is the input activation and **u**(*t*) is the instantaneous input. The time constants of recurrent units and inputs are set by *α*_r_ and *α*_in_. Weights within and between units are housed in the matricies **W**_EE_, **W**_EI_, **W**_IE_, **W**_II_. Only the excitatory units receive projections from the input and project to the output through **W**^in^ and **W**^out^, respectively.

RNNs received two input streams **u**(*t*) = [*u*_1_(*t*), *u*_2_(*t*)] representing sensory evidence:

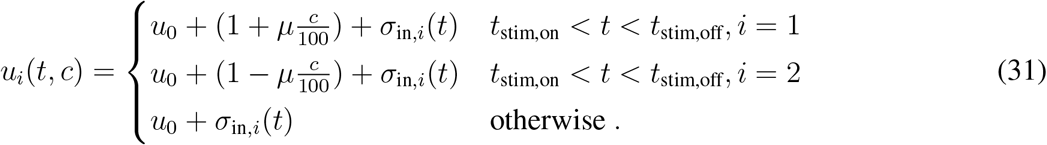

The stimulus period is 21 time steps and *t*_stim,on_ and *t*_stim,off_ are uniquely chosen for each trial. The stimulus magnitude, *μ* = 3.2, is fixed and stimulus difficulty is set by *c* which can range between −20 and 20.

The recurrent and input noise are modeled by the elements of 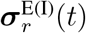 and ***σ***_in_(*t*) that are sampled from a Gaussian distribution. We ensure that each element has a standard deviation *σ*_0,r_ and *σ*_0,in_ via scaling:

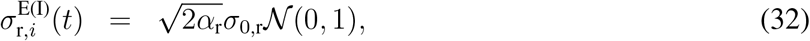

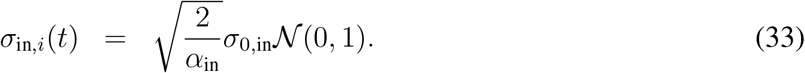

### RNN training

The goal of RNN training is to minimize the difference between the output **z** (*N*_trial_ × *N*_time_ × *N*_out_) and targets **T** (*N*_trial_ × *N*_time_ × *N*_out_). We set the entries in **T** to the baseline value of 0.2 and, following a stimulus onset, raise the entries to 1 for the output corresponding to the correct choice. This target is designed to train the network to remain in a low activity state until stimulated and elevate the correct output in response to a stimulus. Half of training trails (*f*_catch_ = 0.5) were catch trials, on which no stimulus was presented and target values remained at 0.2 throughout the trial. The training batch consisted of *N*_trial_ = 200 trials which were randomly generated every training epoch. Within the training batch, noncatch trials are equally divided between possible choices and the difficulty is randomly sampled.

Recurrent network weights were randomly initialized from a Gamma distribution with a shape *w_μ_* = 0.0375 and scale *w_σ_* = 0.5 for excitatory weights **W**^EE^,**W**^EI^, and *γw_μ_* and scale *w_σ_* for inhibitory weights **W**^IE^, **W^II^**. *γ* = *N*_E_*s*_E_/*N*_I_*s*_I_ scales the strength of inhibitory connections to offset for differences in the number and excitability between excitatory and inhibitory units. Input and output weights **W**^in^, **W**^out^ were randomly initialized from a uniform distribution and then values were normalized so the weights associated each input and output summed to 1 across units. All weights were trained via back propagation through time to minimize the loss function:

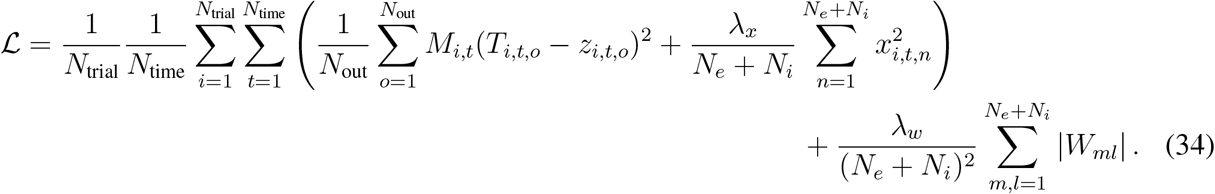

Here **x** is a concatenation of **x**_*E*_ and **x**_*I*_ (*N*_trial_ × *N*_time_ × (*N*_E_ + *N*_I_)), and **W** is a concatenation of **W**^*EE*^, **W**^*EI*^, **W**^*IE*^, and **W**^*II*^ (*N*_E_ × *N*_I_). To encourage the network to integrate the stimulus for extended time, we used a mask **M** (*N*_trial_ × *N*_time)_, where entries were zero during the stimulus period so that time points during the stimulus were not considered when calculating the error term of the loss function. On catch trials, all entries of **M** were set to 1. The hyperparameter λ*_x_* = 0.1 controls the amount of L2 regularization intended to minimize the activation of each unit. The hyperparameter λ*_w_* = 1.0 controls the amount of L1 regularization applied to weights. We updated the weights by stochastic gradient descent using the ADAM optimizer in PyTorch with a learning rate 0.01. During training, the norm of the gradient was clipped at 1.

To maintain the identity of excitatory and inhibitory units and to keep the input and output weights positive, all elements of **W**^*EE*^, **W**^*EI*^, **W**^*IE*^, **W**^*II*^, **W**^in^, and **W**^out^ which are negative were set to 0 after every training step. We prevent self-connections by elementwise multiplying **W**^*EE*^ and **W**^*II*^ by (**1** – **I**), where **I** is the identity matrix and **1** is a matrix of 1s, after every training step.

We terminated RNN training based on its task performance. We tested RNN performance on a validation batch of trials after every training epoch. Each validation batch consisted of 100 trials with stimuli ranging between −20 and 20 in steps of 2. The network registered a decision when the difference between the output variables was above a threshold of 0.25. Trials were considered valid if at least 75% of the prestimulus period was below the decision threshold and at least 50% of the post stimulus period was above the decision threshold. Overall performance was measured as the fraction of correct choices out of all trials except for the ambiguous case where stimulus was equal to 0. We compute the accuracy and the psychometric function only using valid trials. We terminated training when a network’s overall performance reached 85%. RNN parameter values are shown in Table 2.

**Table 2.**
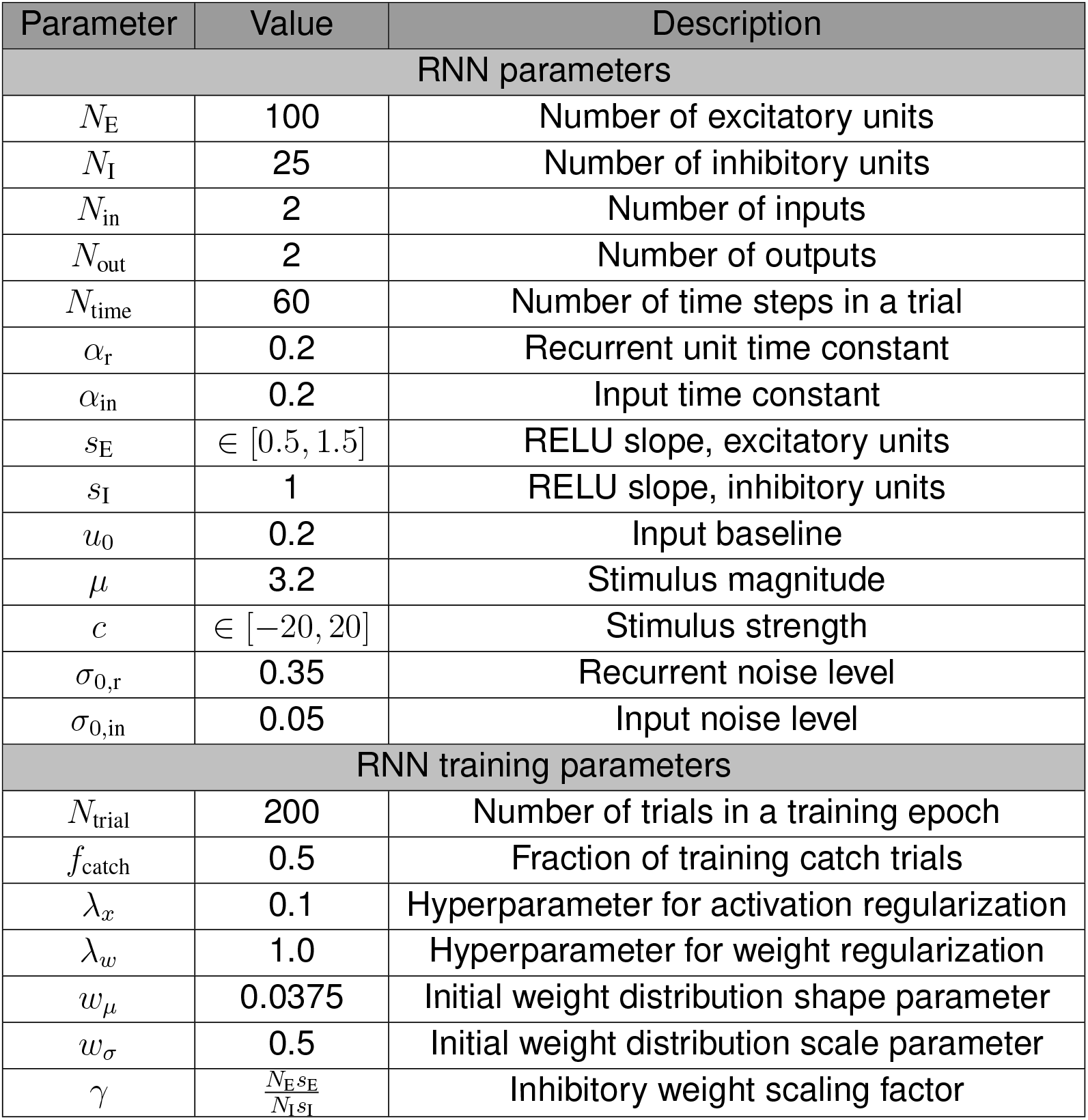
Recurrent neural network parameters.

### Measuring choice selectivity of RNN units

After training, we analyzed the activity of excitatory and inhibitory RNN units to quantify their choice selectivity. Our metric is based on the ability to decode the choice registered by the network based on the activity of the unit at the time point immediately following stimulus offset^21^. For each unit, we computed the receiver operating characteristic (ROC) using the roc function and the area under the ROC curve (AUC_ROC_) using the trapz function in Matlab. A unit with the same activity for either choice will have an AUC_ROC_ equal to 0.5, thus our choice selectivity measure was defined by AUC_ROC_ – 0.5. To identify significantly selective units, we compared AUC_ROC_ to a shuffled distribution generated from that unit’s activity by shuffling the choice outcomes 150 times. We considered units to be choice selective if their AUC_ROC_ fell within the lowest or highest 2.5% percentiles of the shuffled AUC_ROC_ distribution.

### Measuring connection specificity Σ in RNNs

We measured the specificity of connections between choice selective units in RNNs. For each connection class (EE, EI, IE, and II), we computed 〈*w*^+^〉 and 〈*w*^−^〉, the mean strength of the weights between significantly selective units with, respectively, the same and opposite selectivity. Then we computed Σ as:

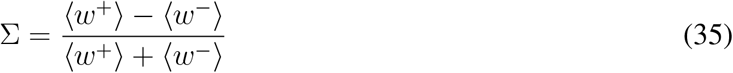

This expression is identical to the Σ used in the mean-field model. To assess significance of correlations between Σ for the 4 connection classes, we computed a shuffled distribution constructed by shuffling the network labels 5,000 times.

### Perturbing inhibitory populations

We perturbed activity of inhibitory neurons by delivering the same constant input to all inhibitory neurons during the stimulus period. In the mean-field model, we modified the parameter *ν*_0, *I*_ by a small amount within the range [–0.5,0.5] around a baseline. We used two baseline values of *ν*_0, *I*_: 11.5 for low-inhibitory regime and 14 for high-inhibitory regime. In RNNs, we delivered perturbations in a similar manner, where we delivered a constant input within the range ∈ [–1,1] during the stimulus period.

## Acknowledgements

This work was supported by the NIH grants F32MH123011 (J.P.R.), R01 EB026949 (A.K.C. and T.A.E.), and 2R01EY022979 (A.K.C.), Alfred P. Sloan Foundation Research Fellowship (T.A.E.), and the ISQEB program at the Simons Center for Quantitative Biology at CSHL (J.P.R.). We thank M. Genkin for thoughtful comments on the manuscript.

## Author contributions

J.P.R., A.K.C and T.A.E. designed the research. J.P.R. developed the code and performed computer simulations. J.P.R., A.K.C and T.A.E. wrote the paper.

## Data availability

The data used in this study can be reproduced using the source code.

## Code availability

The source code to reproduce the results of this study will be available on GitHub upon publication.

## Supplementary figures

**Supplementary Figure 1.**
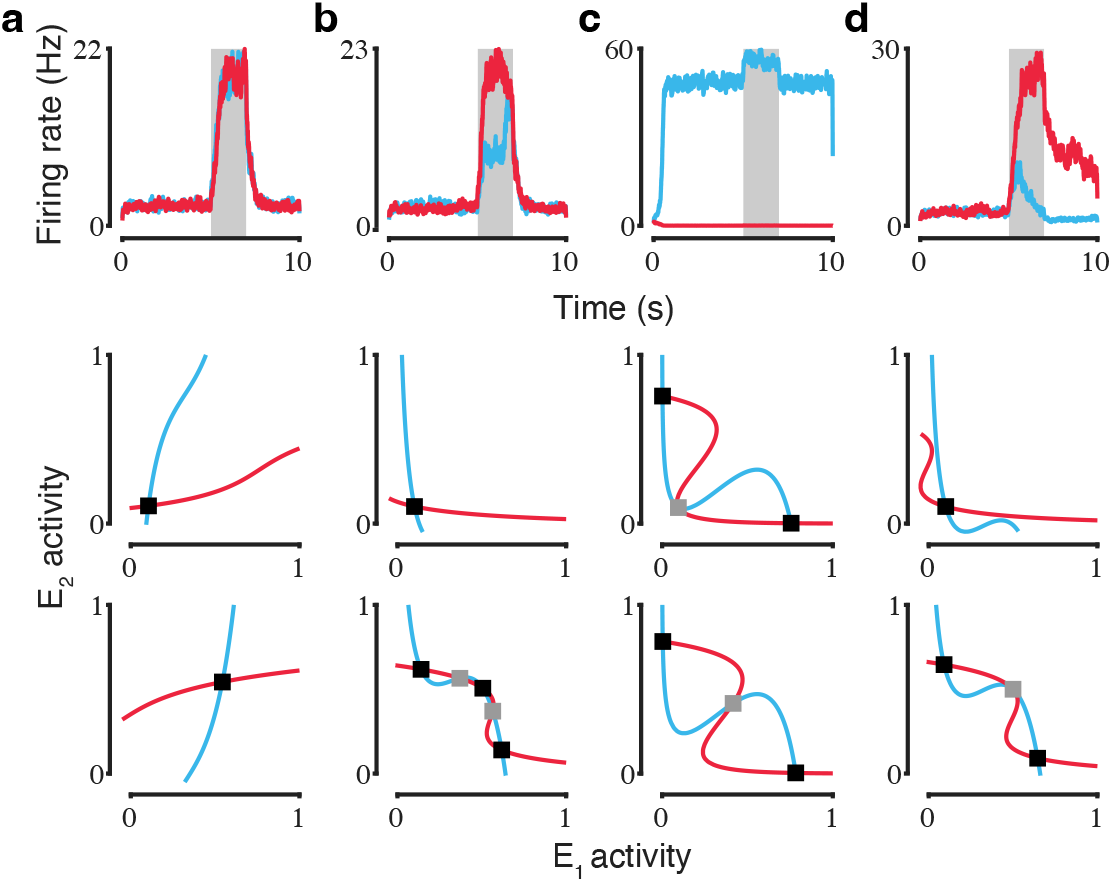
Examples of circuits not capable of decision-making. Connection specificity between choice selective populations determines which fixed points are present in the circuit. Example activity traces for choice selective excitatory populations (upper row, gray area indicates stimulation period with 0 stimulus strength) and the phase plane for unstimulated (middle row) and stimulated circuits (lower row, 0 stimulus strength) are shown. Black and grey squares indicate stable attractors and saddle points, respectively. (**a**) A circuit which lacks choice and working memory attractors, as well as a symmetrical saddle point when stimulated. Σ^EE^ = 0.175, Σ^EI^ = 0, Σ^IE^ = 0. (**b**) A circuit which lacks working memory attractors and the symmetrical saddle point. Σ^EE^ = 0.175, Σ^EI^ = −0.675, Σ^IE^ = 0.675. (**c**) A circuit which lacks the symmetrical low activity attractor. Σ^EE^ = 0.475, Σ^EI^ = 0, Σ^IE^ = 0. (**d**) A circuit which lacks working memory attractors. Σ^EE^ = 0.35, Σ^EI^ = 0.5, Σ^IE^ = 0.45.

**Supplementary Figure 2.**
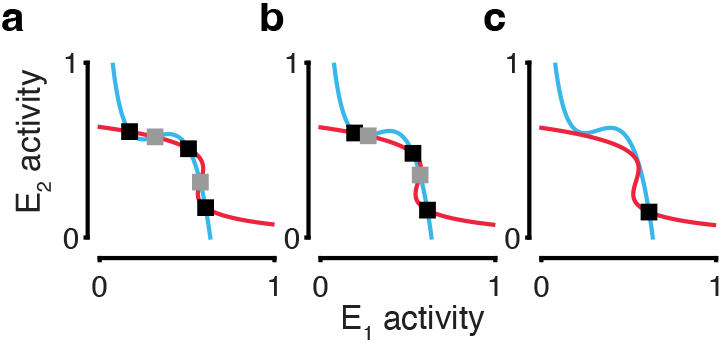
Circuits lacking the saddle point can discriminate easy stimuli. Phase planes of a strongly ipsispecific circuit (Σ^EE^ = 0.3196, Σ^EI^ = 0.25, Σ^IE^ = 0.75) show that as the stimulus strengths toward one of the choices increases, the symmetrical attractor disappears enabling the circuit to make decisions. (**a**) Stimulus strength is 0.0. (**b**) Stimulus strength is 2.5. (**c**) Stimulus strength is 5.0.

**Supplementary Figure 3.**
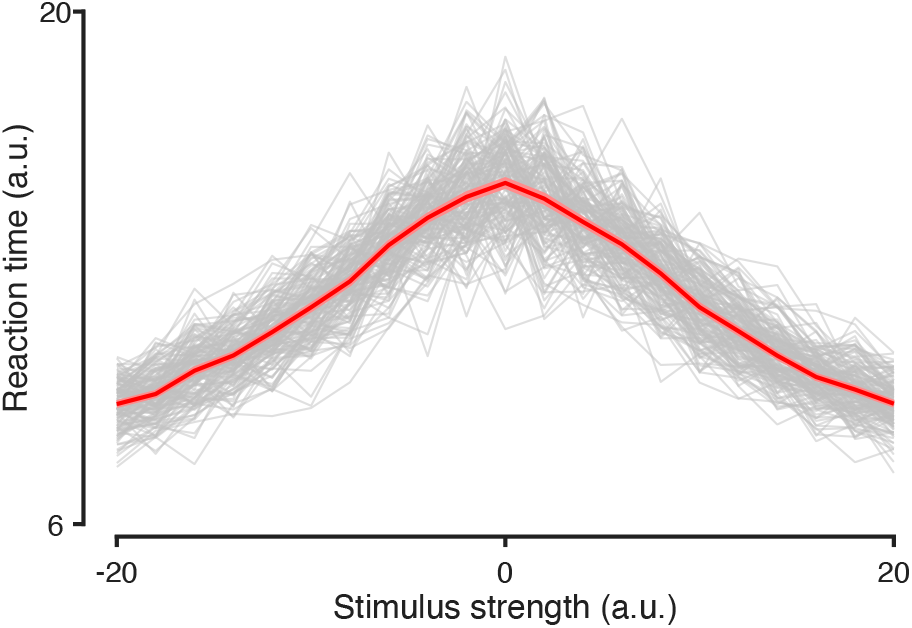
RNNs display a stimulus-strength dependent speed versus accuracy tradeoff. Chronometric functions for individual trained RNNs (grey) and their average (red). RNNs take longer to report decisions for difficult stimuli (stimulus strength near 0) than for easier stimuli.

**Supplementary Figure 4.**
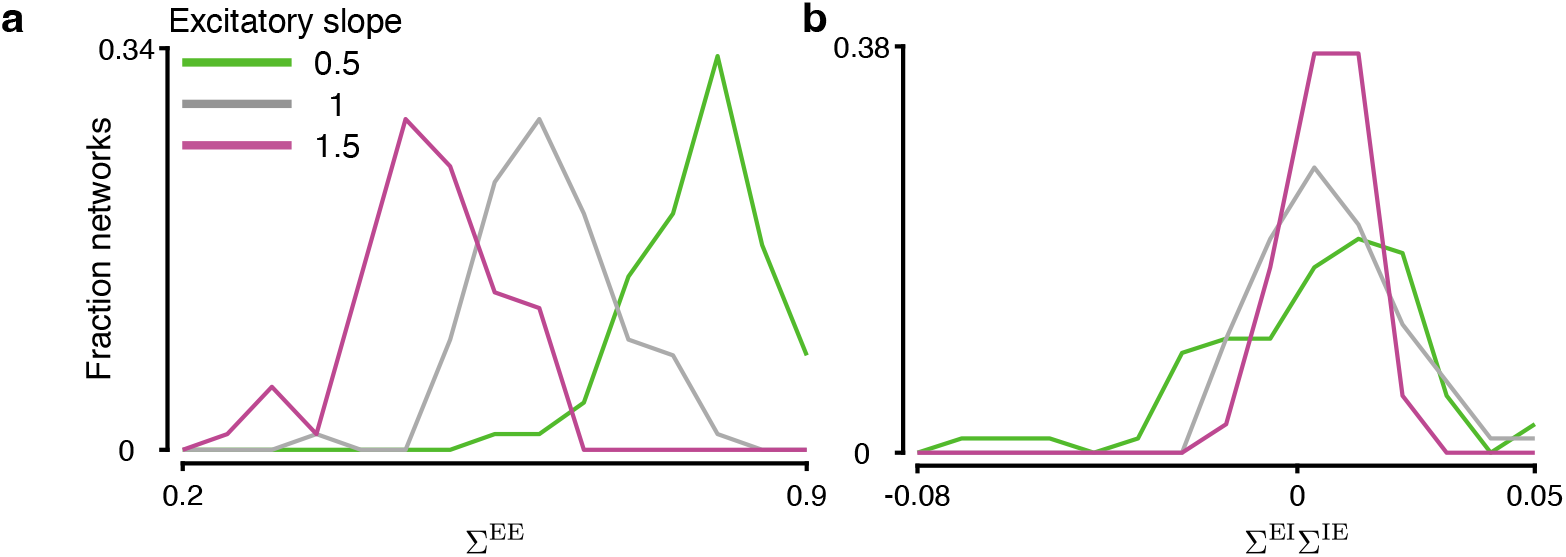
Changes in the excitatory unit excitability result in altered circuit structure in RNNs. (**a**) Trained RNNs with hypoexcitable excitatory units (green) show higher Σ^EE^ than RNNs with baseline (gray) or hyperexcitabile units (purple). (**b**) The distribution of the selectivity index Σ^EI^Σ^IE^ is similar across all RNNs. In all panels, distributions include 75 networks for each excitability level.

**Supplementary Figure 5.**
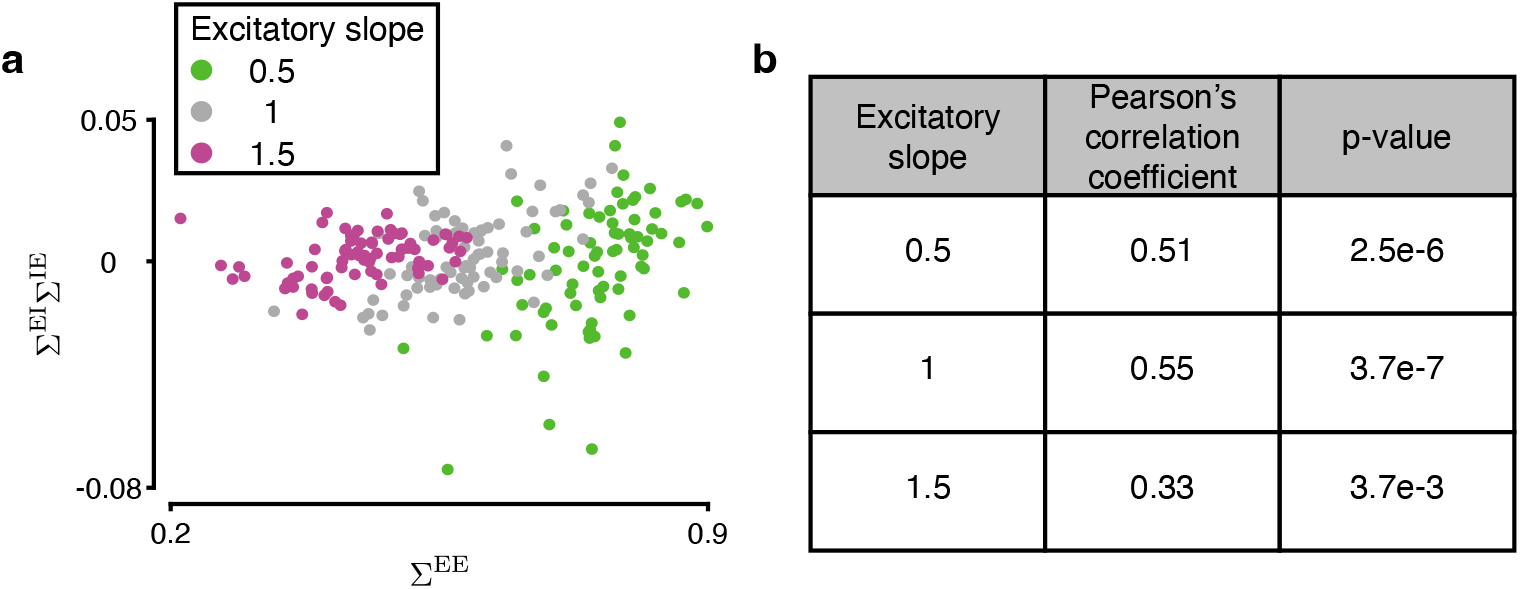
Excitatory and inhibitory selectivity is correlated in RNNs with different excitability of excitatory units. Σ^EE^ and Σ^EI^Σ^EI^ are correlated in RNNs trained with different excitability of excitatory units. (**a**) Hypo- and hyperexcitable networks show a similar relationship between Σ^EE^ and Σ^EI^Σ^IE^ as the baseline networks. (**b**) For each excitability level, the correlation between Σ^EE^ and Σ^EI^Σ^IE^ was significant.

**Supplementary Figure 6.**
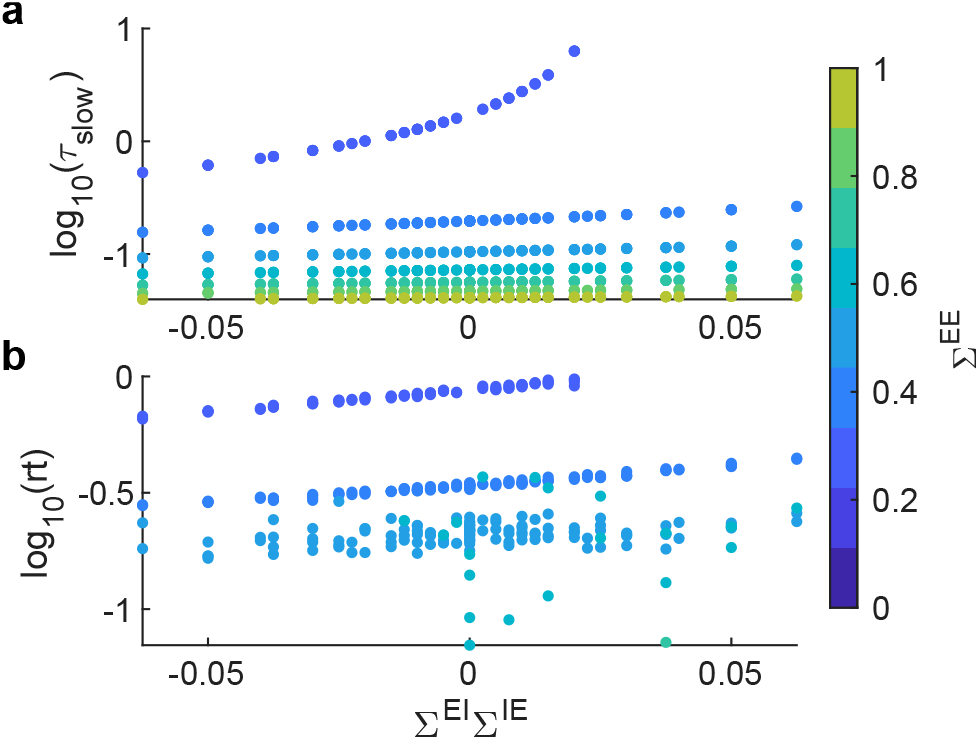
Excitatory selectivity has a larger effect on mean-field circuit dynamics than inhibitory selectivity. Changes in Σ^EE^ (colormap) have a larger effect on *τ*_slow_ (**a**) and reaction time (**b**) than inhibitory selectivity (x-axis).

